# An integrated single-cell RNA-seq map of human neuroblastoma tumors and preclinical models uncovers divergent mesenchymal-like gene expression programs

**DOI:** 10.1101/2023.04.13.536639

**Authors:** Richard H. Chapple, Xueying Liu, Sivaraman Natarajan, Margaret I.M. Alexander, Yuna Kim, Anand G. Patel, Christy W. LaFlamme, Min Pan, William C. Wright, Hyeong-Min Lee, Yinwen Zhang, Meifen Lu, Selene C. Koo, Courtney Long, John Harper, Chandra Savage, Melissa D. Johnson, Thomas Confer, Walter J. Akers, Michael A. Dyer, Heather Sheppard, John Easton, Paul Geeleher

## Abstract

Neuroblastoma is a common pediatric cancer, where preclinical studies suggest that a mesenchymal-like gene expression program contributes to chemotherapy resistance. However, clinical outcomes remain poor, implying we need a better understanding of the relationship between patient tumor heterogeneity and preclinical models. Here, we generated single-cell RNA-seq maps of neuroblastoma cell lines, patient-derived xenograft models (PDX), and a genetically engineered mouse model (GEMM). We developed an unsupervised machine learning approach (‘automatic consensus nonnegative matrix factorization’ (acNMF)) to compare the gene expression programs found in preclinical models to a large cohort of patient tumors. We confirmed a weakly expressed, mesenchymal-like program in otherwise adrenergic cancer cells in some pre-treated high-risk patient tumors, but this appears distinct from the presumptive drug-resistance mesenchymal programs evident in cell lines. Surprisingly however, this weak-mesenchymal-like program was maintained in PDX and could be chemotherapy-induced in our GEMM after only 24 hours, suggesting an uncharacterized therapy-escape mechanism. Collectively, our findings improve the understanding of how neuroblastoma patient tumor heterogeneity is reflected in preclinical models, provides a comprehensive integrated resource, and a generalizable set of computational methodologies for the joint analysis of clinical and pre-clinical single-cell RNA-seq datasets.

## BACKGROUND

Neuroblastoma is a neuroendocrine tumor, typically arising in the adrenal gland, and is the most common extracranial solid tumor in children^1^. Survival in high-risk neuroblastoma is approximately 50%, with survivors suffering long-term consequences of chemotherapy^2^. This is despite new therapies showing clear promise in cell lines and mouse models^3^, indicating that we have an incomplete understanding of how neuroblastoma patient tumor heterogeneity is reflected in preclinical models.

Pediatric tumors, including neuroblastoma, exhibit very few recurrent somatic mutations compared to adult tumors^4^ and heterogeneity at the level of cell state^5^ is thought to contribute to therapy resistance^6–10^. For example, it was recently shown that many neuroblastoma cell lines exist in an admixed state, expressing two discrete super-enhancer-driven gene expression programs, referred to as the “*adrenergic*” (expressing sympathoadrenal features) and the “*mesenchymal*” (comparatively chemoresistant) programs^6,10^. These programs exhibited mutually exclusive expression, where cells expressing one program could interconvert to the other *in vitro*^11,12^.

Single-cell RNA-seq represents the ideal tool to study this non-genetic tumor heterogeneity. Indeed, in the past three years, at least seven different studies^13–19^ have reported single-cell RNA-seq maps of neuroblastoma patient tumors. However, there were disagreements among some of these studies, e.g., with multiple different cell types posited as the neuroblastoma cell-of-origin (chromaffin cell^16^, sympathoblast/neuroblast^15,17,19^, Schwann cell precursors^13^, “sympatho-and chromaffin cells”^18^ and “a subtype of TRKB+ cholinergic progenitor”)^18,15,17–19^. Additionally, arguments were made both in favor of^13–15,18^ and against^16,17,19^ the *in vivo* relevance of the cell-line-derived,^6,10^ putatively-chemoresistant mesenchymal-like gene expression program. Thus, the gene expression programs contributing to therapy resistance in neuroblastoma—and the preclinical models that can be used to study these—remains unclear.

Here, we generated single-cell RNA-seq data from neuroblastoma cell lines, PDXs, and genetic mouse models, using these data to create the first integrated map of human neuroblastoma patient tumors and preclinical models. To achieve this, we developed a novel computational method called “automatic consensus nonnegative matrix factorization” (acNMF; details below). Using acNMF, we determined that the dominant adrenergic gene expression programs were preserved across human tumors and preclinical models. In contrast, the presumptive drug-resistant mesenchymal program demonstrated less consistent behavior, and while clearly expressed in cell lines, exhibited high expression primarily in cancer-associated normal cells *in vivo*. Notably, we confirmed a distinct weak-mesenchymal program in predominantly adrenergic cells of some high-risk patient tumors, which we showed could be elicited 24 hours after administering chemotherapy *in vivo*. Overall, our findings provide a generalizable set of computational tools that we have used to improve the understanding of neuroblastoma patient tumor heterogeneity and preclinical models, which will help inform drug development for this catastrophic disease.

## RESULTS

### Automatic consensus non-negative matrix factorization (acNMF) can accurately recover gene expression programs in single-cell RNA-seq data

NMF is an unsupervised learning approach often used to estimate gene expression programs in single-cell RNA-seq datasets. Using this method, each program is represented as a vector of weights (or “loadings”) assigned to each gene (see, e.g., Fig. 3e). The activity of each program is estimated in individual cells, scoring high if genes with high weights are highly expressed in that cell (see, e.g., Fig. 3g). Like other methods, NMF can discover discrete cell types, but also, potentially, biological processes and cell state programs—factors believed to influence therapy resistance in cancers including neuroblastoma. However, there is no existing method to determine *a priori* the number of gene expression programs in a single-cell RNA-seq dataset, which can lead to inconsistent NMF-based representations of similar data. This makes it difficult to compare tumors and preclinical models across multiple datasets. To overcome this, we first aimed to create a method for identifying the number of gene expression programs (typically called “*k*”) in our datasets. We eventually developed a generalizable method called acNMF (Fig. 1a).

**Figure 1:**
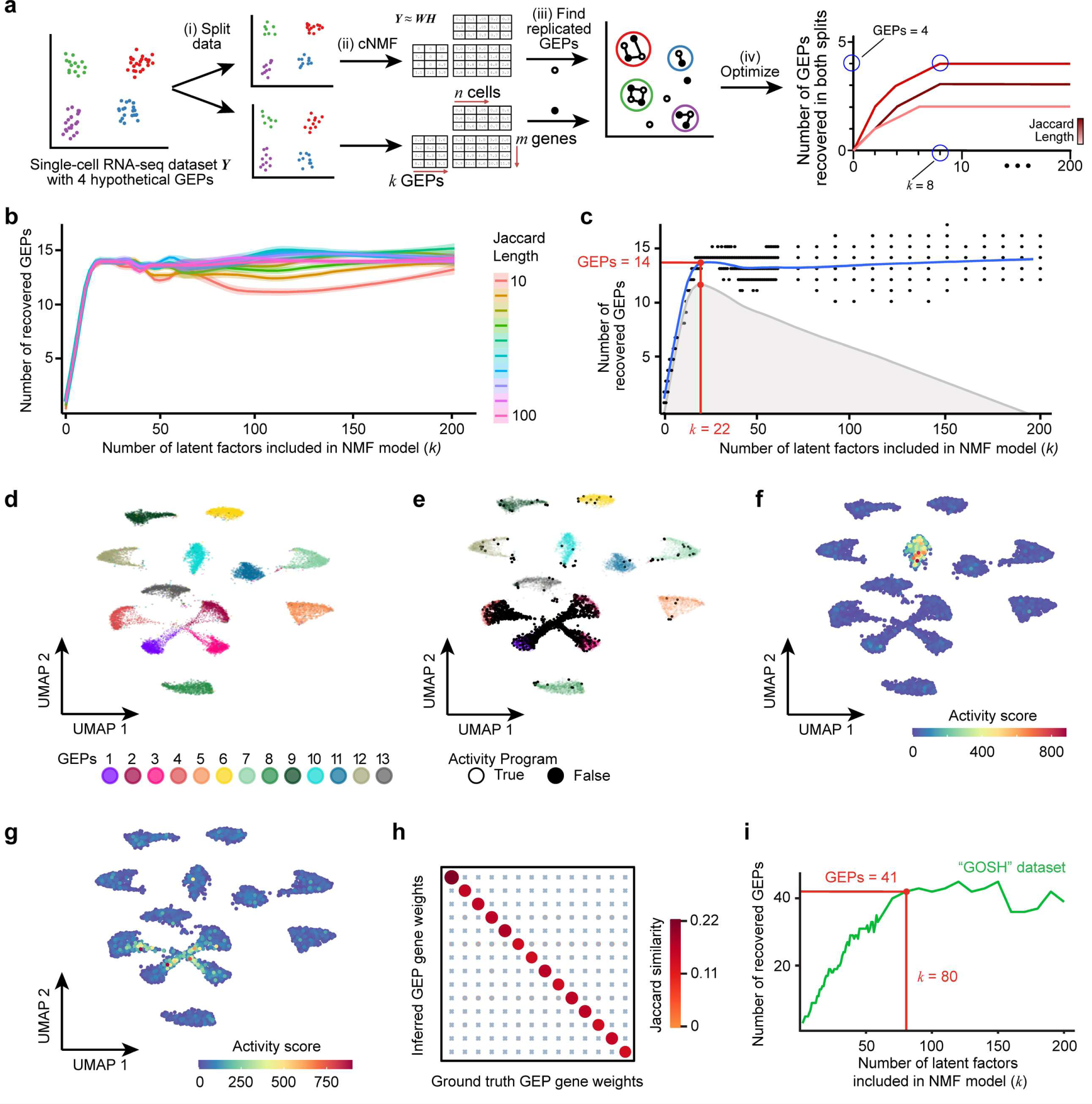
Overview of acNMF methodology and performance in simulated data. (a) Schematic representation of acNMF approach for recovering gene expression programs in a single-cell RNA-seq dataset denoted ***Y***. The dataset is (i) randomly split 50/50, then (ii) each split is factorized independently into a basis matrix of gene expression programs (GEPs) ***W*** and a GEP activity matrix ***H*** using cNMF at a range of values of *k* (e.g. 5-200). At each *k* the redundant GEPs from each split are (iii) reaggregated by a community detection algorithm. This communit number is then (iv) determined for a range of values of *k* and a range of Jaccard length values, parameters which are chosen to maximize the number of GEP-communities replicated in each split. Critically, the number of ground truth GEPs is typically smaller than the value of *k* (GEPs < *k*, hypothetical example highlighted in blue). (b) The number of GEPs reproducibly recovered in both splits of the simulated dataset (y axis) plotted against the value of *k* from NMF (x axis), over a range of values of the Jaccard length (color scale). The approach is repeated 200 times for each pair of values of *k* and Jaccard length and a loess regression curve is fit through these points, with 95% confidence intervals shown. (c) Optimal rank *k* (x axis) is determined by calculating the inflection point (i.e., maximum residual value (gray profile)) of smoothed data in (b) (blue line). (d) UMAP plot of simulated scRNA-seq data, where each dot is a cell colored by the thirteen ground-truth simulated cell types. (e) Like (d), but with black dots highlighting cells expressing the activity program, which (akin to a biological process/pathway) is expressed across multiple simulated cell types. (f) UMAP plot showing a representative example of a recovered ground truth cell identity GEP by acNMF in the simulated single cell RNA-seq dataset. Dots are colored by the activity score of the recovered gene expression program. (g) Like (f) but showing the successful recovery of the activity program. Similar plots for all other recovered programs are shown in Supplementary Fig. 1e-r. (h) Dot plot comparing Jaccard similarity index values of 14 inferred acNMF GEPs with the 14 ground truth GEPs after multiple testing correction (blue X’s represent non-significant comparisons). Both the size and color of the points are scaled by Jaccard similarity. (i) Like (b), but for the GOSH dataset. Data shown for a Jaccard length of 20, which was optimized using acNMF. 41 GEPs were reproducibly recovered in independent splits of the data, at a *k* = 80 (highlighted by the red lines). Similar plots for all datasets in this paper are shown in Supplementary Fig. 1t. GEP; Gene expression program.

Briefly (see Methods for complete details), we initially developed acNMF using a simulated single-cell RNA-seq dataset with 13 cell types and one shared “activity program” (simulating e.g. a biological process). Our inspiration came from observing how the existing cNMF algorithm^20^ behaved on this dataset. We noticed that the 14 ground truth gene expression programs could be identified at most values of *k*, with other programs fitting noise (Fig. 1b). By plotting the value of *k* against the number of reproducibly identified gene expression programs in independent splits of the data, we could find an inflection point (Fig. 1c, Supplementary Figs. 1a-d) where all ground truth programs were recovered and noise-filtered (Fig. 1d-h; Supplementary Figs. 1e-r). In the simulated data, the 14 ground truth gene expression programs were recovered at a *k* value of 22. At *k*=14, only 13 of the 14 programs were recovered (Supplementary Fig. 1s, u). This finding implies that recovering ground truth gene expression programs using NMF-based methods should be decoupled from selecting the appropriate *k*. This is an important general consideration when applying NMF to single-cell RNA-seq data, challenging the idea that finding an optimal *k* value necessarily recovers *k* biologically meaningful programs. We used this method to estimate gene expression programs throughout this manuscript (Fig 1i, Supplementary Fig. 1t; see subsequent sections).

### The acNMF approach produces an interpretable representation of a human neuroblastoma single-cell RNA-seq dataset

Given the range of existing conclusions in the neuroblastoma literature, we decided to first establish an initial baseline set of gene expression programs in patient tumors before comparisons against preclinical models. To that end, we first applied acNMF to the published “GOSH” dataset from Kildisiute *et al.*^17^. The five samples therein cover low-to-high risk neuroblastoma and included one sample from normal adrenal gland. Previously, the insights derived from this dataset were confined to the identification of six cell types. However, the acNMF approach identified 41 gene expression programs (at *k* = 80; Fig. 1i), suggesting possible new insights. Two co-authors (Y.K. and P.G.) blindly and independently annotated these 41 programs. This was done using a web-based platform we developed, presenting each annotator with a catalogue of relevant information, including the dominant gene weights (loadings) for each program, program-program co-expression patterns, inferred copy number profiles, gene ontology term enrichments, cell type annotations from SingleR^21^, and 314 neuroblastoma-relevant marker gene sets that we curated from the literature^10,13–17,19,22–29^ (Supplementary Table S1). The web-based platform presented to annotators is available at http://pscb.stjude.org.

The two annotators assigned qualitatively similar annotations to 37 out of 41 programs (Fig. 2a; Supplementary Table S2). The annotations contained four broad groups of cell types, neuroblastoma cancer cells, immune cells, stromal cells, and endothelial cells (Fig. 2a). We also identified seven “activity” programs that captured processes such as the cell cycle and antigen presentation by MHC. A total of 32 discrete cell types or cell states were recovered, with seven assigned to the stromal group, three to endothelial, 19 to immune, and three to neuroblastoma cancer cells, aided by inferred copy number data (Supplementary Fig. 2a-f; see Methods; further details in subsequent subsections). We refer to these three neuroblastoma-specific programs as “Adrenergic I (sympathoblast-like)”, “Adrenergic II (pre-neuronal-like)”, and “Neuroblastoma-MYC” (NB-MYC). All three neuroblastoma programs were strongly enriched for the classical cell line derived adrenergic marker genes (Fig. 3a-c; 8.1 × 10^-66^ for Adrenergic I, *P* = 1.7 × 10^-72^ Adrenergic II, 1.1 × 10^-23^ for Neuroblastoma-MYC from one-sided Wilcoxon rank sum test, signature defined by Van Groningen *et al.*^10^), but consistent with a subset of previous studies^16,17,19^, were not strongly enriched for mesenchymal markers.

**Figure 2:**
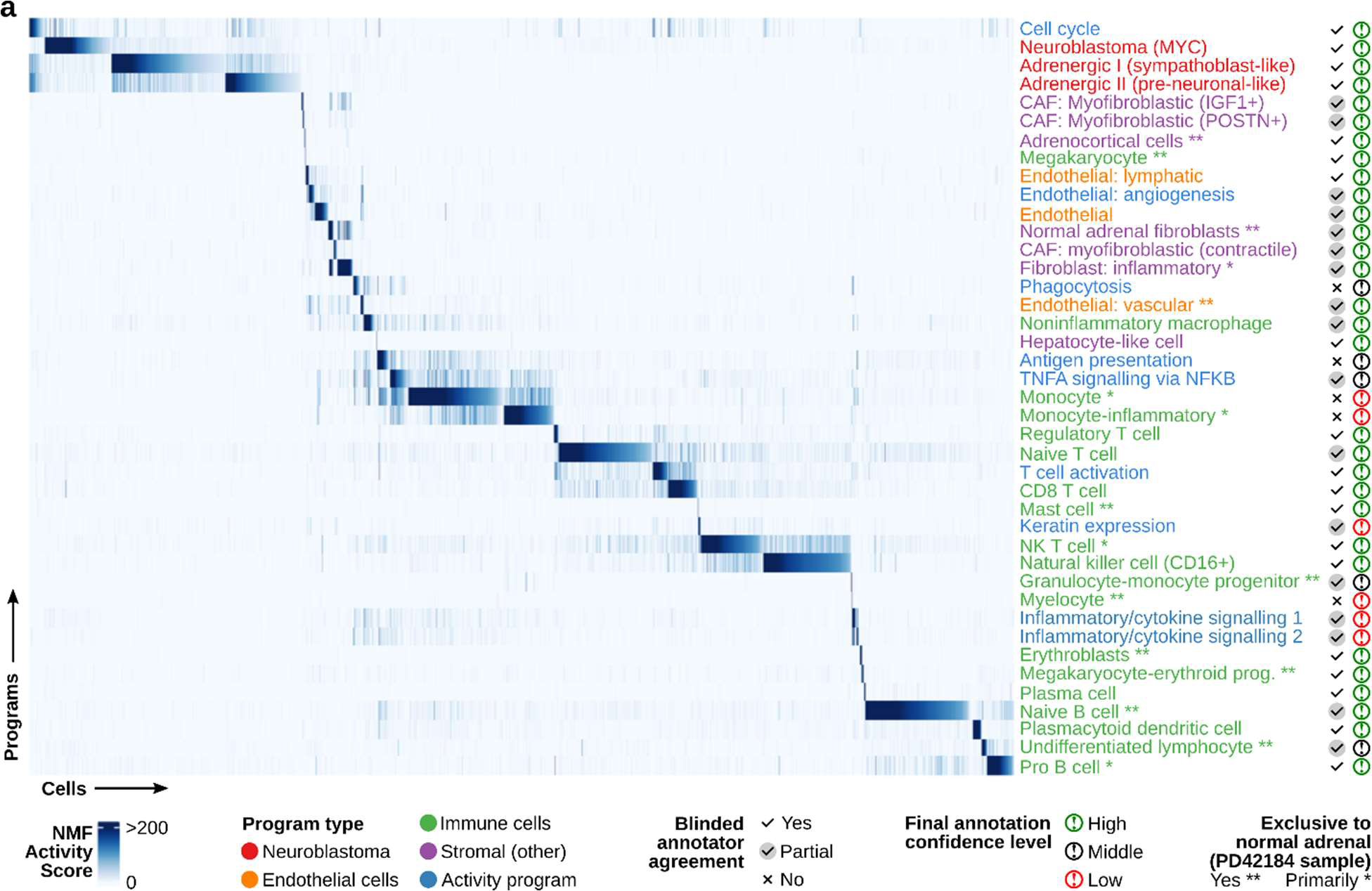
Gene expression programs recovered by acNMF in the GOSH neuroblastoma single-cell RNA-seq dataset. (a) Heatmap showing the activity scores (color scale) for the 41 recovered programs (y axis) across each single cell (x axis) in the GOSH dataset. The final consensus annotation for each gene expression program is shown on the right, colored by the type of program. Tick marks indicate whether the two blinded annotators agreed or not, and the far-right column indicates whether the annotators reached a high, middle, or low level of confidence in the final annotation.

**Figure 3:**
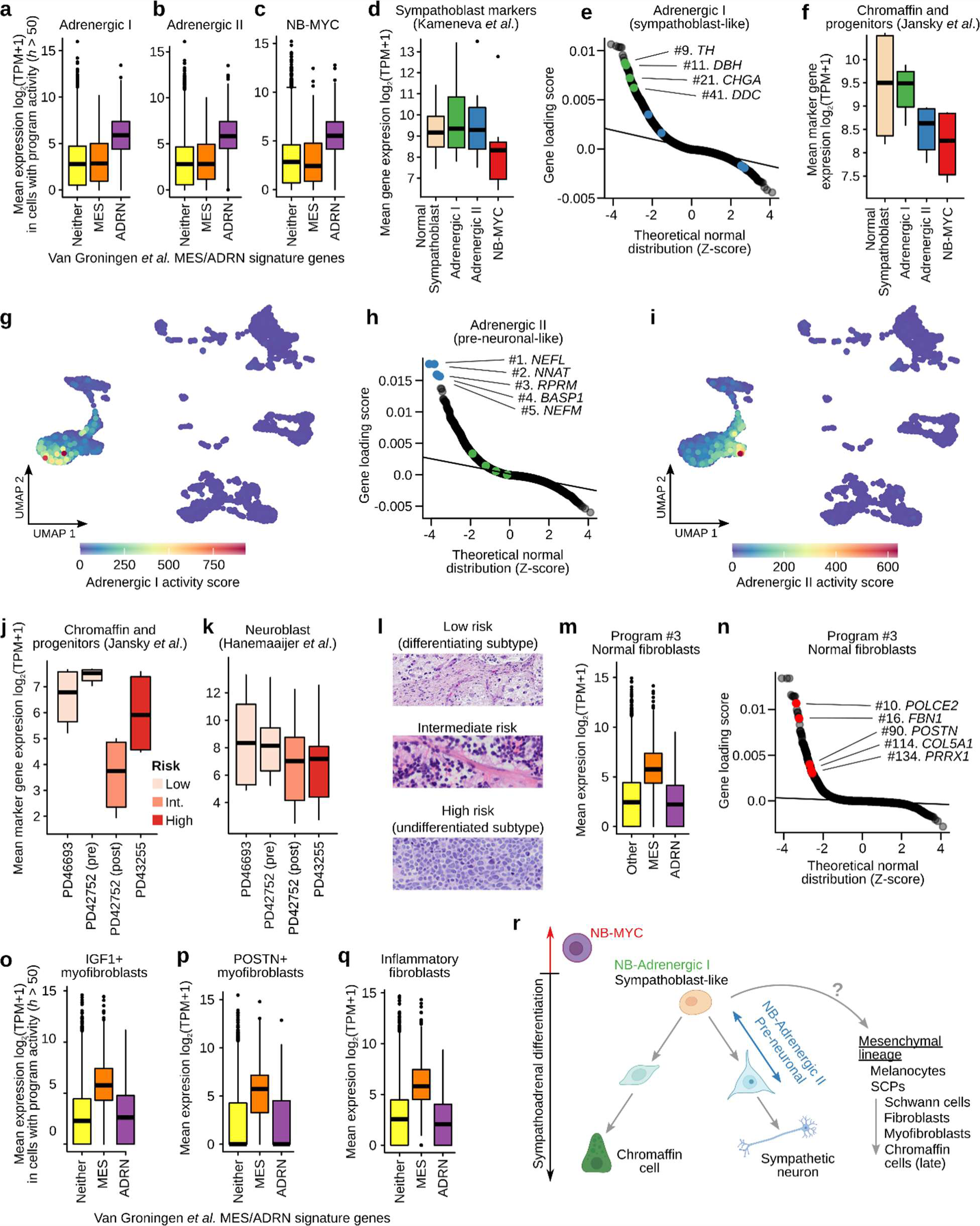
Cells expressing multiple adrenergic and mesenchymal-like gene expression programs are evident in neuroblastoma tumors. (a) Boxplots showing the mean expression (y axis) of the genes from the classical Van Groningen mesenchymal (MES; orange box) or adrenergic (ADRM; purple box) signature, in cells that are highly expressing (activity *h* > 50) the acNMF-identified “Adrenergic I (sympathoblast-like)” program in the GOSH dataset. The yellow box shows the mean expression of all other genes in these cells. (b) Like (a) but for the “Adrenergic II (pre-neuronal-like)” program. (c) Like (a) but for the “Neuroblastoma-MYC” program. (d) Boxplots showing the mean gene expression (y axis) of sympathoblast marker genes (curated from Kameneva *et al.*) in cells highly expressing (activity *h* > 50) each of the 3 acNMF recovered adrenergic programs, or in normal sympathoblasts (obtained from the Descartes Cell Atlas). (e) QQ-plot showing the gene weights (loading scores) (y axis) for the “Adrenergic I (sympathoblast-like)” program. Key sympathoblast marker genes have been highlighted in green, and key neuronal marker genes in blue. (f) Like (d) but for the “Chromaffin and progenitors” marker gene set, curated from Jansky *et al*.| (g) UMAP plot of the GOSH dataset; each point represents a single-cell, colored by the activity of the “Adrenergic I (sympathoblast-like)” program. (h) Like (e) but for the “Adrenergic II (pre-neuronal-like)” program. (i) Like (g) but for the “Adrenergic II (pre-neuronal-like)” program. (j) Boxplot showing the mean gene expression of “Chromaffin and progenitors” marker genes (curated from Jansky *et al*.) in the cancer cells of the 4 neuroblastoma samples profiled in the GOSH dataset. Boxes are colored by tumor risk classification. (k) Like (j) but for the “Neuroblast” marker gene set curated from Hanemaaijer *et al*. (l) Representative H&E stains of low, intermediate and high-risk human neuroblastoma tumors, illustrating varying degrees of differentiated features. (m) Boxplots showing the mean expression (y axis) of the genes from the classical Van Groningen mesenchymal (MES; orange box) or adrenergic (ADRM; purple box) signature, in cells that are highly expressing (activity *h* > 50) the acNMF-identified “Normal fibroblasts” program in the GOSH dataset. The yellow box shows the mean expression of all other genes in these cells. (n) QQ-plot showing the gene loading scores (y axis) for the “Normal fibroblasts” program. A subset of genes classically used as markers of mesenchymal-like neuroblastoma cells are highlighted in red. (o) Like (m) but for the “IGF1+ myofibroblasts” program. (p) Like (m) but for the “POSTN+ myofibroblasts” program. (q) Like (m) but for the “Inflammatory fibroblasts” program. (r) A schematic diagram overlaying the acNMF-recovered programs on normal sympathoadrenal differentiation^34^.

### The dominant neuroblastoma adrenergic programs resemble distinct aspects of sympathoadrenal development

For all three adrenergic programs, the top ranked curated human cell-type signature enrichment was sympathoblast (Fig. 3d, *P* = 1 × 10^-6^, 3.6 × 10^-3^, 7.6 × 10^-6^ for adrenergic I, adrenergic II, NB-MYC respectively, marker genes curated from Kameneva *et al.*^15^). Sympathoblasts are a neural crest-derived precursor associated with the development of the sympathetic nervous system and this result is consistent with a subset of the existing neuroblastoma single-cell RNA-seq studies^15,17,19^. However, acNMF uncovered additional complexity beyond a clean resemblance to sympathoblasts: Specifically, the Adrenergic I (sympathoblast-like) program was defined by the expression of genes involved in the synthesis of norepinephrine in both sympathoblast-precursors and differentiating chromaffin cells (Fig. 3e-g). This includes tyrosine hydroxylase (*TH*), dopamine beta hydroxylase (*DBH*), dopa decarboxylase (*DDC*) (“Chromaffin and progenitors” gene list curated from Jansky *et al.*^19^). Notably, markers specific to mature chromaffin cells (*PNMT* and *PENK*) were not enriched, consistent with the argument that neuroblastoma cells do not typically resemble differentiated chromaffin cells^26,30,31^.

The Adrenergic II (pre-neuronal-like) program over-expressed markers of neuroblasts defined by Hanemaaijer *et al*.^24^ and Jansky *et al.*^19^.The top five highly expressed marker genes included neuronal-specific genes, neurofilament light chain (*NEFL*), neuronatin (*NNAT*), brain acid soluble protein 1 (*BASP1*), and neurofilament medium chain (*NEFM*). This program can be co-expressed with the Adrenergic I program (Fig. 3g & 3i, Supplementary Fig. 3a-c). Thus, these programs were independent but not mutually exclusive.

The third recovered neuroblastoma program, NB-MYC, was only expressed in the high-risk patient sample (PD43255) and was characterized by lower expression of marker genes of differentiating chromaffin and neuronal cell types (Fig. 3d,3f, Supplementary Fig. 3a), consistent with the undifferentiated pathology of high-risk neuroblastoma^32^. The defining genes of this program were dominated by translation initiation and elongation factors (Supplementary Figs. 3d & 3e), consistent with high levels of transcription and translation supporting the hyper-aggressive growth of high-risk tumors. The top marker genes of the adrenergic I & II programs were expressed 2-3-fold lower in the high-risk tumor (Fig. 3j-k, *P* = 6.9 × 10^-3^ and *P* = 5.9 × 10^-2^ respectively). This is consistent with low/intermediate-risk tumors exhibiting differentiating histology (Fig. 3l)^32^. Adrenergic marker genes were also strongly differentially expressed between high-risk tumors and other tumors in 498 bulk RNA-seq samples^33^ (Supplementary Fig. S3f-h; *P* = 1.8 × 10^-16^, 1.8 × 10^-5^, and 1.1 × 10^-18^ for Adrenergic I, II and MYC programs respectively), suggesting these are generalizable patterns of differential gene expression.

While our study does not aim to definitively pinpoint neuroblastoma’s cell-of-origin, these results are consistent with the proposition that neuroblastoma tumors broadly resemble adrenergic neuroblasts/sympathoblasts^15,17,19^. However, we additionally propose that these tumors can co-express a mixture of independent adrenergic subprograms, resembling subtly different sympathoadrenal cell types (Fig. 3g, i, Supplementary Fig. 3b-c), and that these activities differ between high and low/intermediate risk disease. This suggests that the adrenergic co-expression patterns in neuroblastoma patient tumors are more nuanced than previously described. Evidence for these behaviors in other datasets and preclinical models is presented in a later subsection.

### Cancer associated fibroblasts subpopulations in the neuroblastoma tumor microenvironment strongly express mesenchymal features, heavily resembling mesenchymal-neuroblastoma cell lines

We identified nine gene expression programs in the GOSH dataset that statistically resembled the neuroblastoma cell line-derived presumptive-drug-resistant mesenchymal gene expression program. This includes some very strong enrichments (e.g., *P* = 1.78 × 10^-88^ for Program 40 for enrichment of Van Groningen “mesenchymal” signature, Supplementary Table S2). This seems to imply that mesenchymal neuroblastoma cells have been identified in these tumors. However, one of these programs was expressed exclusively in fibroblasts of the reference normal adrenal sample (Fig. 3m-n; *P* = 3.75 × 10^-40^ for mesenchymal enrichment) and thus this specific program cannot be expressed in cancer cells. Hence, these shared gene expression features of fibroblasts and mesenchymal-like neuroblastoma cell lines suggest a source of disagreement about whether mesenchymal-like neuroblastoma cells exist *in vivo*. Indeed, in the four neuroblastoma tumors, our acNMF decomposition again identified several programs strongly resembling mesenchymal-like neuroblastoma cell lines. However, in all cases, we annotated these as non-cancerous fibroblast cell populations rather than as mesenchymal-like neuroblastoma cells (Fig. 3o-q; these annotations leveraged inferred copy number - details in subsequent section). Notably, however, acNMF revealed a much more complex landscape of cancer-associated fibroblasts (CAFs) than previously reported in neuroblastoma^28^. Specifically, we identified IGF1+ (*P* = 1.92 × 10^-7^), POSTN+ (*P* = 3 × 10^-15^) and contractile myofibroblast populations (*P* = 2.5 × 10^-3^). We also identified inflammatory CAFs (*P* = 2.9 × 10^-9^) expressing interleukin (*IL6*, *IL33*) and complement component genes (*C7*, *C3*) using marker genes from Lavie *et al*.^28^. Overall, our acNMF-based decomposition of the GOSH dataset yielded a conceptual model with which to describe the behavior of neuroblastoma cancer cells in patient tumors (Fig. 3r) but did not identify clear evidence for mesenchymal-like neuroblastoma cancer cells. We next explore whether these initial characteristics are recapitulated in other human datasets and the range of preclinical models.

### The fidelity of neuroblastoma gene expression programs across human tumors and preclinical models

Next, to assess the generality of the gene expression programs found in the GOSH dataset and assess their fidelity in neuroblastoma preclinical models, we assembled five more of the publicly available patient tumor datasets: from Dong *et al.*, Jansky *et al.*, Kameneva *et al.*, Verhoeven *et al.*, and the PMC dataset from Kildisuite *et al.*^15–19,35^, totaling 69 samples. Additionally, we performed single-cell RNA-seq profiling across 13 neuroblastoma cell lines, 17 patient derived xenografts (PDX), and 11 samples from a genetic mouse model (GEMM)^36^ (metadata for this (largest-of-its-kind) cohort is assembled in Supplementary Table S3, and in our web platform).

We applied the acNMF decomposition to each of these datasets independently, yielding between 17 (cell lines) and 45 (Dong *et al.*) gene expression programs per dataset (Supplementary Fig. 1t, Supplementary Table S4). To pairwise cross-compare the programs from each dataset, we represented the programs as nodes on a graph. We drew an edge between nodes if the program was replicated in another dataset (*FWER* < 0.01 by an asymmetric Jaccard similarity test; Fig. 4a; Supplementary Fig. 4a; see Methods). A force-directed algorithm facilitated the layout of the graph whereby interconnected nodes are found in closer proximity than sparsely connected nodes. The resulting graph’s structure indicated that similar gene expression programs had often been recovered in multiple datasets, including preclinical models. Consensus hubs (see Methods) were evident for gene expression programs representing several immune cell types, neuroblastoma-specific programs, stromal cells such as fibroblasts, and activity programs such as the cell cycle and hypoxia (Fig. 4a-d).

**Figure 4:**
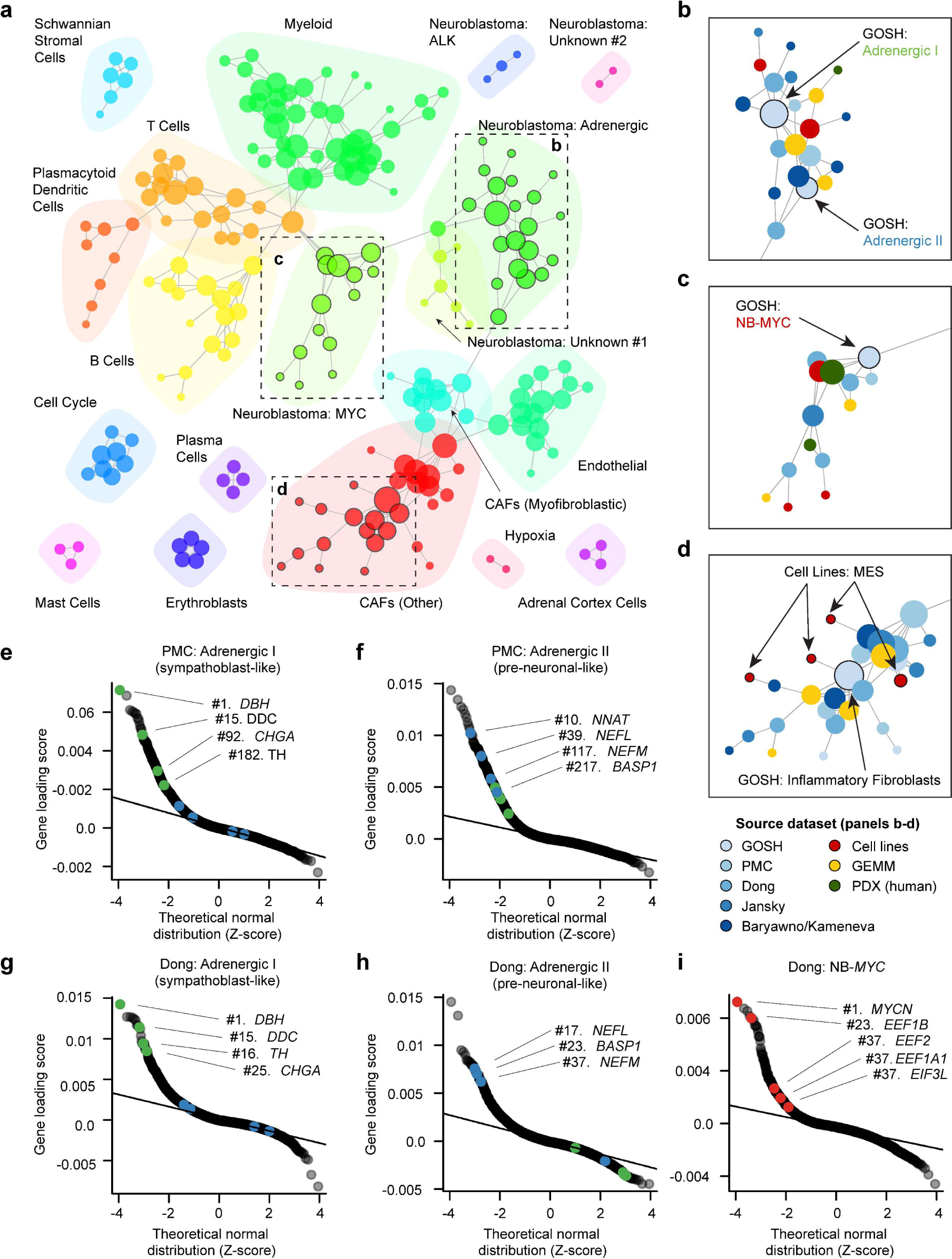
The fidelity of neuroblastoma gene expression programs across human tumors and preclinical models. (a) A graph of gene expression programs identified by acNMF across eight neuroblastoma single-cell RNA-seq datasets, from humans, PDX, GEMM and cell lines. Nodes represent a single program’s gene weights (i.e. loading scores), and node size is determined by the degree (i.e. # of connections). Edges connect similar nodes by a Jaccard similarity test. The colors highlight related communities of nodes. The black boxes highlight regions of interest, which are shown in detail in panels (b-d). (b) Inset showing a detailed view of region (b) from panel (a). This region has connected Adrenergic I and Adrenergic II programs. In this inset, nodes have been colored by dataset, rather than community (as in panel (a)). (c) Like (b) but for a dense interconnected community of “Neuroblastoma MYC” programs. (d) Like (b) but showing a region for a community of gene expression programs expressed in cancer associated fibroblasts. A neuroblastoma program, exclusive to the GIMEN cell line, is connected to this community of nodes. (e) QQ-plot showing the gene weights (loading scores) (y axis) for the “Adrenergic I (sympathoblast-like)” program recovered in the PMC dataset. Similar to Fig. 3e from the GOSH dataset key sympathoblast marker genes have been highlighted in green, and key neuronal marker genes in blue. (f) Like (e) but for the “Adrenergic II (pre-neuronal-like)” program. (g) Like (e) but for program 24 “Adrenergic I (sympathoblast-like)” in the Dong *et al.* dataset. (h) Like (e) but for program 19 “Adrenergic II (pre-neuronal-like)” in the Dong *et al.* dataset. (i) Like (e) but showing the “Neuroblastoma MYC” program from Dong *et al*. Top marker genes, similar to those identified in the GOSH dataset, have been highlighted in red. Note: An interactive version of the graph plot (panel a), and QQ-plots (panels e-i) for every program in every dataset are available in our web platform.

The adrenergic I (sympathoblast-like) program from GOSH was directly or indirectly connected to programs in all other datasets, including all preclinical models (Fig. 4b). These connected programs retained the characteristic hallmarks, including overexpression of genes like *DBH*, *TH* and *CHGA* (Fig. 4e-h). The adrenergic II pre-neuronal-like program was also observed in multiple other datasets (Fig. 4f & h) and the “neuroblastoma-MYC” (high risk) program formed a dense hub (Fig. 4c), with similar programs recovered in high-risk (both *MYC* and *MYCN* overexpressing) tumors in all datasets (Fig. 4i).

This cross-dataset analysis also revealed three groups of novel programs expressed in neuroblastoma cancer cells. We were able to find a potential explanation for one, which was expressed in *ALK* activated tumors (Fig. 4a; supported by one patient tumor and two PDXs but no other preclinical model). Additionally, we found five programs unique to a single human neuroblastoma tumor, implying a possible long tail of divergent neuroblastoma programs that currently remain unexplained and are absent in our profiled preclinical models.

The mesenchymal-like programs observed in the GOSH dataset formed a large mesenchymal/fibroblast subgraph. Interestingly, programs from cancer cell lines were also connected to this fibroblast subgraph. For example, a program expressed exclusively in the GIMEN cell line^10^ was directly connected to the inflammatory fibroblasts program in the GOSH dataset (Fig. 4d), again consistent with similar gene expression features in mesenchymal-like neuroblastoma cells in culture and untransformed fibroblasts *in vivo* (Fig. 3o-q).

Overall, this cross-dataset-integrated map argues that, despite some disagreements in the literature, neuroblastoma single-cell RNA-seq datasets are reasonably consistent and the dominant adrenergic programs are preserved in all preclinical models. However, the status of mesenchymal-like tumor cells and cell lines requires further exploration. Note: this graph-based representation of these datasets (Fig. 4a) is available as part of our interactive, web-based platform (accessible at http://pscb.stjude.org).

### Cells highly expressing a mesenchymal-like program are consistent with non-cancerous cells in all datasets except cell lines

Our cross-dataset analysis above also revealed other candidate mesenchymal-like neuroblastoma cells, including Schwann-like cells (e.g., Dong dataset, Program 27, *P* = 1 × 10^-9^ for enrichment of Descartes Schwann cell markers, *P* = 5.3 × 10^-43^ for Van Groningen mesenchymal enrichment; Supplementary Fig. 5a). We were interested in whether any of these could plausibly represent *in vivo* mesenchymal neuroblastoma cancer cells.

We therefore applied InferCNV^37^, a computational method for inferring copy number variant (CNV) profiles from single-cell RNA-seq data, across the entire integrated largest-of-its-kind cohort (see Methods). We found that the CNV profiles of mesenchymal-like cells were always similar to the reference normal, whereas CNV aberrations in adrenergic cells were typically obvious (Supplementary Fig. 2, Supplementary Figs. 5a-c). The exception was cell lines, where both mesenchymal and adrenergic cells always exhibited massive CNV aberrations consistent with cancer cells (Supplementary Fig. 5d-f).

However, because CNV-based inference tools act at the level of clustered groups of cells, InferCNV cannot rule out a small proportion of mesenchymal-like cancer cells. We therefore constructed a machine learning model that, by leveraging the inferred copy number profiles of the high-confidence-cancer and high-confidence-normal cells, could *predict* the cancer/normal status of the ambiguous mesenchymal-like cells on a per-cell basis (Supplementary Fig. 5g; Supplementary Tables 5 & 6; see Methods). These models could predict the cancer/normal status of the held-out cancer/normal-endothelial cells with median accuracy approaching 100% (Fig. 5a, Supplementary Fig. 5h). We applied the trained models to the held-out ambiguous population of mesenchymal-like cells (candidate mesenchymal neuroblastoma cells, cancer-associated fibroblasts, or Schwann-like cells). The model never classified mesenchymal cells as cancer cells beyond the model’s misclassification rate estimated from normal endothelial cells (Fig. 5b-c; Supplementary Fig. 5i-n). Thus, these analyses, given the scale of the data currently available, did not provide convincing evidence for subsets of neuroblastoma cells that highly express a mesenchymal-like gene expression program *in vivo*.

**Figure 5:**
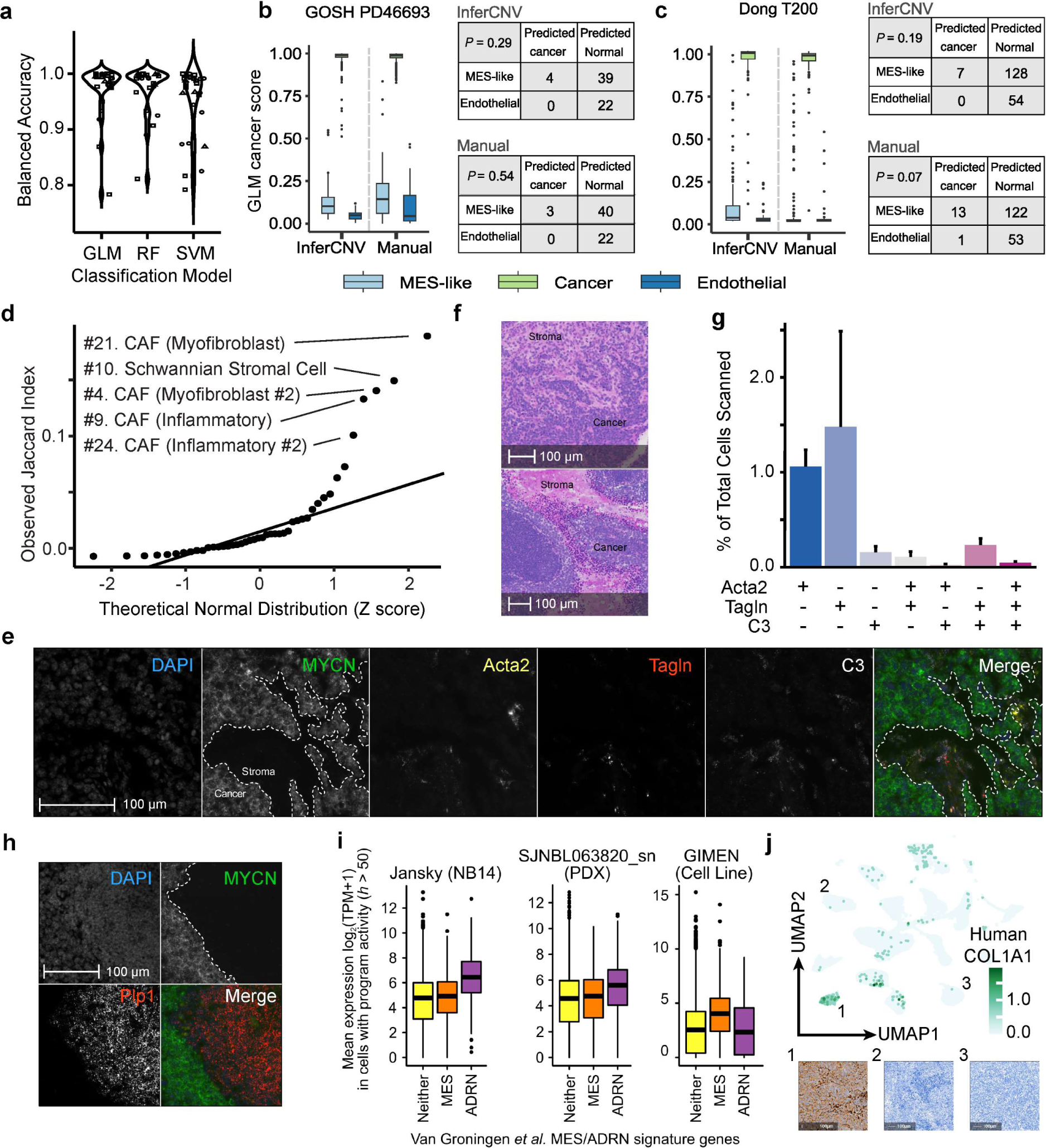
The cancer/normal status of ambiguous mesenchymal-like cells in neuroblastoma tumors. a) Violin plots comparing the balanced accuracy (y axis) for prediction of cancer/normal status of known cancer or immune cells for the 3 different initial models (x axis) across 13 different tumor samples (shown for both “InferCNV” and “manual” features (see methods)). GLM, generalized linear model; RF, random forest; SVM, support vector machine. b) Boxplots showing the predicted probability of each cell being a cancer cell from the GLM model (y axis) for the ambiguous-mesenchymal cells (light blue), for high confidence cancer cells held out from the initial model fitting (green) and for normal endothelial cells held out from the initial model fitting (dark blue). The predictions are shown for models using features defined based on InferCNV (left 3 boxes) or from manual bins of 50 adjacent genes (right 3 boxes). The contingency tables (right panels) estimate whether the number of mesenchymal-like cells classified as cancer cells differs statistically from the misclassification rate of normal endothelial cells, calculated by a Fisher’s exact test. This panel shows a representative sample PD46693 from the GOSH dataset. c) Like (b) but for the Dong sample T200, all other samples are in Supplementary Fig. 5 and 8. d) QQ-plot showing the Jaccard Similarity (y axis) of other programs with Program #37 (Myofibroblastic CAF) in our GEMM dataset. e) Representative RNA ISH image of CAF markers in GEMM tumors. Neuroblastoma markers (human *MYCN,* and *Ddc*) were used to define cancer cells. DAPI counterstaining confirmed the presence of cells located in regions that lack *MYCN* expression. The co-expression of CAF markers was identified within the *MYCN*-absent stromal regions (NB831 shown). f) H&E staining of mouse tumors identifies regions of densely packed dark stained neuroblastoma cells, and pink-staining stromal cells in our GEMM, which resemble the *MYCN*-absent regions of mesenchymal marker expression in our RNA ISH experiments. g) Bar plot showing the co-expression of CAF markers in GEMM tumors (y axis). All combinations of CAF marker expression were quantified in stromal cells (x axis), and they are represented as proportions of the total cell numbers identified in each sample. h) Representative image of GEMM tumors stained with *MYCN* and the Schwann cell marker *Plp1* in NB839. *Plp1* expression was restricted to *MYCN*-absent regions in these tumors. i) Boxplot showing the mean expression (y axis) of genes in neuroblastoma cancer cells from the sample NB14 from Jansky *et al*. Boxes are genes from the classical Van Groningen mesenchymal (MES; orange box) or adrenergic (ADRN; purple box) signatures, with the yellow box showing the expression of all other genes. The middle panel shows our PDX sample SJNBL063820 and the right panels shows the predominantly mesenchymal neuroblastoma cell line GIMEN. j) Immunohistochemical staining for the mesenchymal-marker gene COL1A1 in our SJNBL063820 PDX sample (#1, left) and in two other PDX samples SJNBL046145 (#2, middle) and SJNBL047443 (#3, right). The upper panel shows a UMAP plot colored by the expression of *COL1A1* in our PDX cohort, with the numbers indicating the UMAP clusters corresponding to the 3 stained PDX samples.

We validated these putative fibroblast and Schwann-like cells using an orthogonal assay—a highly sensitive spatial transcriptomics assay, multiplexed RNA *in situ* hybridization (RNA ISH) (Fig 5d, Supplementary Fig. 6a-c). We applied this in our GEMM tumors, a model that develops neuroblastoma because of Cre-conditional activation of the *human* copy of *MYCN* in dopamine beta hydroxylase (*Dbh*) expressing cells—thus, human *MYCN* expression should be excluded from non-cancer cells. In all tumors, the human *MYCN* transcript was highly expressed across most of the tissue, but absent from some large regions (Fig. 5e, Supplementary Fig. 7a-d) that resemble Schwannian stroma (Fig. 5f, Supplementary Fig. 7e). These *MYCN*-absent regions clearly co-expressed the myofibroblast markers *Tagln* and *Acta2* (Fig. 5e), consistent with the idea that these mesenchymal cells are fibroblasts and not cancer cells. Regions excluding *MYCN* also expressed the inflammatory fibroblast marker *C3* (Fig. 5e). Interestingly, we identified some cells that seemed to co-express all combinations of chosen fibroblast markers, consistent with the notion of intermediate subpopulations (Fig 5g). Schwann cell markers *Sox10* and *Plp1* were also excluded from regions expressing human *MYCN* (Fig. 5h; Supplementary Fig. 7e). Collectively, the data indicate that, despite strong transcriptional resemblance to neuroblastoma mesenchymal cell lines, these *in vivo* cells are predominantly Schwann-like cells and previously uncharacterized fibroblast subpopulations (see Discussion).

### Rare high-risk neuroblastoma tumors express a weakly enriched mesenchymal-like signature *in vivo* that can be recapitulated in PDX

Interestingly, we identified some drug pretreated tumors that seem to represent a partial exception to the above. In high-risk sample NB14 from the Jansky *et al.* dataset, the tumor’s cancer cells were broadly (but weakly) statistically enriched for the Van Groningen cell-line-derived mesenchymal gene expression signature (Fig 5i; Program 12 *P* = 1.7 × 10^-6^; Supplementary Fig. 8a-b). Notably, these cells were still predominantly adrenergic (Fig. 5i, purple boxes). This is different from the behavior of cell lines, where the expression of mesenchymal-like genes corresponds to a loss of expression of adrenergic genes^6,10^ (Fig. 5i).

We initially suspected that this was artefactual, but, surprisingly, we also identified a high-risk pretreated human tumor in our PDX cohort that exhibited a similar weak enrichment of the mesenchymal signature across most cancer cells of the tumor (Fig. 5i; Program 1, *P* = 1.6 × 10^-12^ in SJNBL063820, Supplementary Fig. 8c-d). We obtained this PDX tissue^38^ and used immunohistochemistry to stain for COL1A1 protein, a classical neuroblastoma mesenchymal marker^10^. Remarkably, greater than 95% of the neoplastic cells of SJNBL063820 expressed COL1A1 in the cytoplasm and membrane, but staining was not evident in two negative control PDXs (Fig. 5j, Supplementary Fig 8g). Thus, this weak-mesenchymal phenotype can be recapitulated in PDX. Interestingly, two treatment-naïve high-risk *MYCN* amplified tumors (T200, T230) from the Dong dataset, while still adrenergic (Supplementary Figs. 8e-f), also showed broad overexpression of the transcription factor *TWIST1*, a known driver of epithelial-to-mesenchymal transition^39^ (ranked #9 on Program 11 (T200-specific) and #27 on Program 33 (T230-specific)). Curiously, though, both programs were also weakly but significantly co-expressed with the mesenchymal-fibroblast Program 7 (*r_s_* = 0.07 *P* = 1.4 × 10^-33^ and *r_s_* = 0.06, *P* = 7.1 × 10^-25^ respectively), also consistent with a very weak or proto-mesenchymal cell state expressed in otherwise adrenergic cells. In general, these findings uncover a complex landscape of mesenchymal-like gene expression programs *in vivo*, where the behaviors appear different to cell line models (see Discussion) but can be maintained in PDX.

### A mesenchymal-like gene expression program can be chemotherapy induced in neuroblastoma cells *in vivo*

The confirmation of proto-mesenchymal features in some drug-pretreated tumors in multiple datasets (e.g., NB14, SJNBL063820)—but not in baseline tumors—led us to hypothesize that the weak mesenchymal-like programs could have been chemotherapy induced. To test this, we treated five of our GEMMs with a single pharmacologically relevant dose of cisplatin (a mainstay of neuroblastoma treatment) and treated six GEMMs with vehicle-control (see Methods). We harvested these tumors 24 hours after treatment and performed single-cell RNA-seq (see Methods; Supplementary Fig. 9a).

Consistent with a treatment effect, the drug-treated samples clustered separately from the vehicle-treated samples (Fig. 6a). To determine which genes had been induced/repressed, we created “pseudo bulk” expression profiles using all of the cells of each tumor and then performed functional enrichment (drug vs. vehicle) using GSEA, testing the mSigDB “Hallmark” gene sets (Fig. 6b-d) (see Methods). As expected, “Apoptosis” was among the top chemotherapy-induced gene sets (Fig. 6c-d and 6i, *P* = 0.03; Supplementary Table 7). Remarkably, induced genes were also highly enriched for “Epithelial to Mesenchymal Transition (EMT)” (Fig. 6c-d and 6g, *P* = 7.4 × 10^-3^). We probed this further using acNMF, where in the drug treated samples, we identified (for the first time) clear mesenchymal-like programs expressed in cancer cells (Fig. 6e). Additionally, cisplatin-treated samples were the first where our machine learning model classified cells highly expressing a mesenchymal program as cancer cells (*P* < 2.2 × 10^-16^ for NB853, *P* = 9.5 × 10^-11^ for NB847, Supplementary Figs. 9b-f).

**Figure 6:**
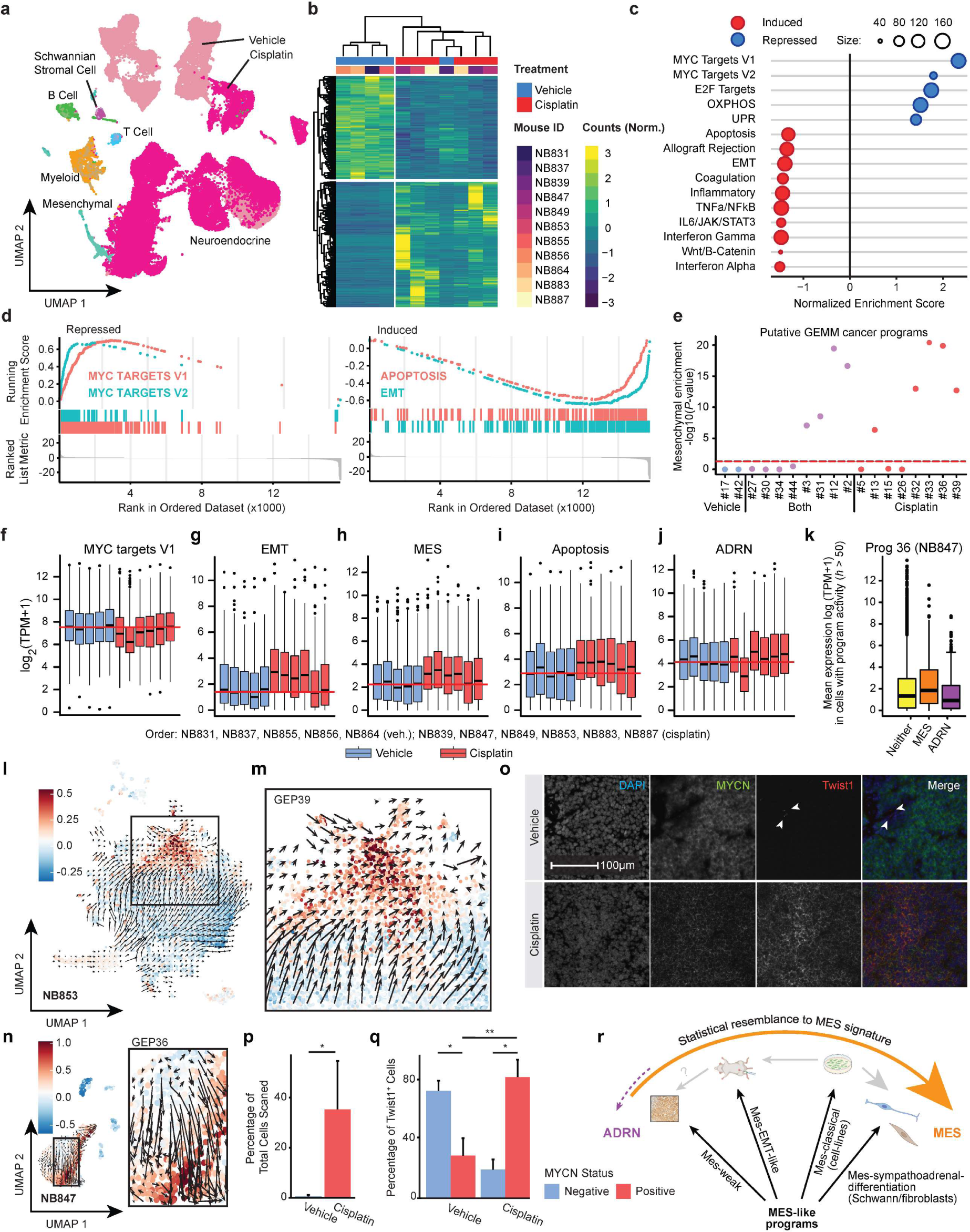
An acutely chemotherapy treated neuroblastoma genetic mouse model strongly overexpresses a mesenchymal-like program in cancer cells. a) UMAP plot of single-cell RNA-seq data obtained from the tumors of all 11 mice. Each point represents a single cell, colored by whether it was obtained from a cisplatin-treated or a vehicle-treated mouse. Brighter colored chroma indicates cisplatin treated animals. b) Heatmap of normalized gene expression values highlighting differentially expressed genes identified using DESeq2. c) Dot plot showing GSEA enrichment scores for the top mSigDB “Hallmark” gene sets. Expression in pseudobulk cancer cells was compared between (*n*=6) cisplatin treated mice and (*n*=5) vehicle treated mice. A negative enrichment score implies the gene set was more highly expressed in drug treated mice. d) GSEA plots summarizing key induced and repressed gene sets. The colored tick marks show the rank of the genes in the gene set of interest and the colored curves show the running enrichment score. e) Statistical enrichment of Van Groningen mesenchymal (MES) signature (y axis) in gene expression programs (x axis) expressed in high-confidence cancer cells in our GEMM. *P*-values were calculated using a Wilcoxon rank sum test (rank of MES genes vs all other genes). f) Boxplot showing the pseudobulk expression in cancer cells from all 11 mice for genes in the Hallmark gene set “MYC targets V1”. Order of samples is NB831, NB837, NB855, NB856, NB864 (vehicle treated, blue); NB839, NB847, NB849, NB853, NB883, NB887 (cisplatin treated, red). The red line indicates the median of vehicle treated samples. g) Like (f) but for Hallmark gene set “Epithelial to mesenchymal transition”. h) Like (f) but for the Van Groningen mesenchymal signature (MES) genes. i) Like (f) but for the Hallmark gene set “Apoptosis”. j) Like (f) but for the Van Groningen adrenergic (ADRN) signature. k) Boxplots showing the mean expression (y axis) of the genes from the classical Van Groningen mesenchymal (MES; orange box) or adrenergic (ADRM; purple box) signature, in cells that are highly expressing (activity *h* > 50) the acNMF-identified program 36, which has characteristics consistent with cancer cells and is expressed primarily in cisplatin treated mouse NB847. The yellow box shows the mean expression of all other genes in these cells. l) UMAP plot showing RNA-velocity analysis of mouse NB853. The cellular trajectory, inferred from splicing dynamics, shown by the arrows, is consistent with cancer cells actively transitioning to a mesenchymal state, which is marked by the expression of the mesenchymal-cancer Program 39 (color scale). m) Detailed view of the box in (l). n) Like (l) but for the sample NB847 and mesenchymal Program 36. o) Representative RNA ISH images for co-localization of *Twist1* with human *MYCN* in vehicle treated (top panels, NB855) and drug treated samples (bottom panels, NB887). p) Bar plot showing the percentage of *Twist1*^+^ cells in each of our sampled mouse slides (y axis), quantified using the HALO software. (*, *P* < 0.05 by t-test, error bars represent SEM) q) Bar plot showing the percentage of *Twist1*^+^ cells (y axis) in either vehicle treated, or cisplatin treated tumors that do or do not co express *MYCN* (colors). (*, *P* < 0.05; **, *P* < 0.01 by t-test) r) Schematic representation of the distinct mesenchymal-like gene expression programs that can be identified across neuroblastoma datasets and preclinical models. These programs have a varying statistical resemblance to the classical cell line derived Van Groningen mesenchymal signature (MES), but despite this, may capture qualitatively different aspects of neuroblastoma cell biology.

While the repressed genes were quite consistent across all drug-treated samples, e.g., the loss of the MYC target genes (Fig. 6b-d and 6f), the patterns of induced genes showed greater variation (Fig. 6b and 6f-j, Supplementary Table 8 for *P*-values). The EMT signal was strongest in NB839, NB847, NB849, and NB853 (Fig. 6g), where the classical Van Groningen mesenchymal signature was also enriched (Fig. 6h). These tumors also had the highest induction of apoptosis genes (Fig. 6i), with a loss of the adrenergic genes in NB847 (Fig. 6j)— reminiscent of trends we had previously observed only in cell lines (Fig. 6k; Fig. 5i). We performed RNA-velocity analysis in NB847 and NB853, the results of which were consistent with the mesenchymal programs resulting from an acute cell state transition (Fig. 6l-n, Supplementary Fig. 10a-e, Supplementary Figure 11a-c). Overall, these recently drug-treated tumors demonstrate that a mesenchymal-like gene expression program can be chemotherapy-induced in neuroblastoma.

Our GEMM, which expresses human *MYCN* in cancer cells, again allowed us to validate these conclusions by an orthogonal RNA ISH assay. This time we assessed the co-localization of human *MYCN* and the classical EMT marker *Twist1* across four drug and four vehicle-treated samples. While *Twist1* expression was evident to some degree in both the drug and vehicle-treated samples, the co-localization of *MYCN* and *Twist1* was much higher in drug-treated samples, again consistent with the induction of a mesenchymal program (Fig. 6o and 6q).

Overall, the landscape of mesenchymal programs in neuroblastoma differs between cell lines and other tumor models. In the *in vivo* setting, this involves weakly enriched expression in still predominantly adrenergic cells (Fig. 6r), which our GEMM data showed could relate to a mesenchymal transition during chemotherapy. Further assessment of the relevance of mesenchymal programs in cell lines to the behaviors of tumors *in vivo* should provide a clear basis for future research (see Discussion).

## DISCUSSION

Multiple preclinical cancer models are commonly applied in drug discovery and mechanistic experimental work. However, few formal methods have been developed to determine how these various models represent/resemble primary patient tumors. Here, we developed a novel NMF-based computational method and a web-based front-end user interface for integrating and interpreting gene expression programs obtained from human tumors and preclinical models. We used these tools to perform the first integrative analysis of single-cell RNA-seq data from human neuroblastoma tumors and preclinical models, and we provide a new public resource for interrogating neuroblastoma biology (http://pscb.stjude.org).

Using our tools, we identified novel neuroblastoma adrenergic programs mimicking the sympathoadrenal developmental lineage and showed that neuroblastoma tumors co-express marker genes of developmental cell types. These behaviors were preserved between human tumors and preclinical models. We propose a conceptual mapping of neuroblastoma gene expression programs to sympathoadrenal development in Fig. 3r, naming these adrenergic programs “Adrenergic I (sympathoblast-like)”, “Adrenergic II (pre-neuronal-like)” and “Neuroblastoma-MYC”.

We also found that the classical cell-line-derived mesenchymal-like program has differences between human tumors and preclinical models. First, while the neuroblastoma mesenchymal signature was clearly identifiable in neuroblastoma cell lines, we could not identify strong evidence that a similar program is expressed in cancer cells in human tumors—neither by inferred copy number nor by RNA ISH. Regardless, it is logically impossible to conclude the non-existence of such cells. Indeed, they could be identifiable in other tumors, by profiling larger numbers of cells, or by using more sensitive technologies. For instance, our RNA ISH experiments have difficulty distinguishing cancer vs. fibroblast/Schwann-like cells at the boundaries of stromal regions, it is conceivable some cell types could have dropped out of the single-cell profiling, and tools for inferring copy number are imperfect^40^. Accordingly, a recent large spatial atlas reported evidence of malignant Schwann-like programs in some neuroblastoma tumors^41^, thus, as the available data continues to grow, it is likely that further nuances will continue to be discovered. However, despite the complexity, we can conclude that mesenchymal-like neuroblastoma cells are likely less abundant than fibroblasts and Schwann-like cells *in vivo*, which also strongly express a similar mesenchymal-like signature and thus, on the basis of marker gene expression, are easily confused with neuroblastoma cells. These analyses also identified novel subpopulations of cancer-associated fibroblasts in most datasets/tumors, which we could validate using RNA ISH in our GEMM, revealing at least inflammatory, myofibroblast, and, potentially, intermediate subpopulations. Similar cell types are critical in regulating immunosuppression, metastasis, and progression in other cancers^28^, and determining the biological/therapeutic relevance of these new cell types in neuroblastoma will form the basis for future research.

Lastly, in multiple datasets we identified rare, high-risk, drug pretreated tumors that weakly express mesenchymal-like features in cancer cells that remain predominantly adrenergic. This contrasts with the behavior in cell lines, where cells expressing a mesenchymal state lose adrenergic features. This may be related to the concept of “high risk tumors with mesenchymal-like features”, which was recently proposed by Jansky *et al*^19^, but in contrast to Jansky we showed these cells at least partially resemble the classical Van Groningen mesenchymal signature and that this could be maintained in PDX. We also identified a potential source of this phenotype, showing that a mesenchymal-like program could be strongly induced *in vivo* in an acutely chemotherapy-treated GEMM. This is consistent with observations across several other cancers that EMT can represent an apoptosis escape mechanism that leads to drug resistance^42,43^. Notably, our observation of the loss of expression of MYC target genes and gain of EMT/mesenchymal-like features in cancer cells also resembles a previously reported “embryonic diapause-like” drug/chemo persistent state change in breast cancer^44^. Our data suggest that high-risk neuroblastoma cells may be primed to undergo a similar transition. It is tempting to speculate that targeting this process could circumvent chemoresistance, and it is plausible that some mesenchymal-like cell lines may represent a good model for this behavior. Because of the very short 24-hour treatment window, this finding also implies that the expression of mesenchymal-like genes need not necessarily emerge in relapsed neuroblastoma tumors by purifying selection, as was previously speculated based on patient data^6,10,45^ and tumors obtained at later time points post-chemotherapy and -relapse in mice^45^. However, the clinical significance of these drug-induced and weakly enriched mesenchymal programs and their potential role in facilitating escape from chemotherapy-induced apoptosis remains to be determined. In general, our results argue that mesenchymal-like programs in neuroblastoma can be subdivided into at least (1) a weak mesenchymal-like signature co-expressed in still-predominantly-adrenergic cells of some high-risk tumors, some of which (2) could potentially be chemotherapy induced; (3) the classical mesenchymal-like signature in cell lines, which also strongly express features similar to (4) cancer-associated fibroblasts and Schwann-like cells found in tumors. As these represent (at least partially) qualitatively different entities, we propose naming these (1) Mes-weak, (2) Mes-EMT-like, (3) Mes-classical, and (4) Mes-sympathoadrenal-differentiation (Fig. 6r).

Finally, our findings only touch on the biological insights that could be gained from an interpretable integrated representation of single-cell RNA-seq data from human neuroblastoma and preclinical models. For example, our NMF-based models recovered a rich representation of the immune component of neuroblastoma tumors, largely recapitulated in our GEMM. We have made all the data and models available in our interactive web-based platform (http://pscb.stjude.org), which we encourage readers to review, discuss, and use as a map for future discoveries.

## CONCLUSIONS

We have developed a set of computational methods for integrative analysis of human single-cell RNA-seq datasets and pre-clinical models. We have used these tools to identify the divergent behavior of gene expression programs in neuroblastoma tumors and preclinical models, providing insight into the behavior of putative drug resistance mesenchymal-like programs. We have provided this dataset as an integrated community resource and our computational approach is applicable in other diseases, where the fidelity of pre-clinical models remains largely unknown.

## METHODS

### Establishing a neuroblastoma genetic mouse model

LSL-MYCN;Dbh-iCre mice^36^ (129X1/SvJ) were obtained from Michael A. Dyer’s lab (St. Jude). This study was carried out in strict accordance with the recommendations of the National Institute of Health’s Guide to Care and Use of Laboratory Animals. The protocol was approved by the Institutional Animal Care and Use Committee at St. Jude Children’s Research Hospital. All mice were housed, per approved IACUC protocols, on a 12−12 light cycle (light on 6 am, off 6 pm) and provided food and water ad libitum. Neuroblastoma-like tumors comparable to human neuroblastoma arose in the abdomen from either the adrenal gland or the celiac and superior cervical ganglia with a penetrance of approximately 50%, at a median age of about 6 months old. The tumor volumes were monitored by ultrasound using the Vevo 3100 system (Fujifilm-Visualsonics, Toronto, Canada), and once an adrenal tumor had reached 500mm^3^ in diameter, the mice were treated with either a single, pharmacologically relevant^46^ 2 mg/kg dose of cisplatin (*n* = 6) or vehicle control (*n* = 5) by intraperitoneal injection. The mice were euthanized at 24 hours, and tumors were immediately harvested and subjected to single-cell RNA-seq using the protocols described below.

### Orthotopic patient-derived xenografts

Flash-frozen fragments of orthotopic patient-derived xenografts were obtained through the St. Jude Childhood Solid Tumor Network (https://cstn.stjude.cloud/)^38,47^. Approximately 50-100 mg of frozen tissue underwent nuclei extraction using manual disruption within a chilled Tween containing buffer^48^. To generate fresh xenograft tissue, cells were injected into the para-adrenal space of immunocompromised nude mice using ultrasound. Tumors were harvested from xenograft-bearing mice, and approximately 250-500 mg of tissue were dissociated using the Papain Dissociation System (Worthington Biochemical)^48^.

### Single-cell/single-nuclear RNA-seq data generation and preprocessing

#### Mouse

Single cells from mouse tumors were obtained through dissociation using the Papain Enzymatic dissociation method following the Worthington kit (cat. No. LK003150), and final dissociated cells were suspended in the DMEM medium.

#### Cell Lines

The following NB cell lines were maintained in the following culture conditions (5% CO_2,_ 37° C, RPMI medium, 10% FBS, 2mM L-glutamine, 100 IU/mL penicillin, 100 μg/mL streptomycin); BE2C, BE2M17, CHP134, CHP212, GIMEN, IMR5, KELLY, MHHNB11, NGP, SKNAS, SKNFI, SKNSH, TGW. Cells were harvested, and viabilty was measured using Lunc Cell Counter AO/PI dye. We processed samples that showed greater than 90% viability.

#### Library Preparation and Sequencing

The dissociated/suspended single cells were filtered and strained through a 40µm strainer and cells were counted using the Luna cell counter. Next, they were loaded into Chromium Chips from Single cell 3’ Gene expression V3.1 kit (10X Genomics, Pleasanton, CA) with target capture of 6000 cells. The cDNA generation, library preparation, and QC was performed following 10X Genomics protocol. The library was sequenced in Novaseq-V1 reagents, and the sequenced data were processed using the Cell Ranger Software (10X Genomics) mapped to mm10 (mouse) or GRCh38 (cell line). Poor quality and outlier cells were identified and removed. Analysis downstream of acNMF, was conducted on normalized read counts (SCTransform, Seurat R package) of the resulting cells.

### Automatic Consensus Non-negative Matrix Factorization (acNMF)

For each dataset, raw read counts were merged into a single matrix containing all samples. 50,000 cells were subsampled per dataset, and then randomly split into two subsets containing roughly equal numbers of cells. cNMF^20^ (https://github.com/dylkot/cNMF/) was applied to each split independently by performing 200 NMF iterations per rank *(k),* spanning a range of *k* = 2 to *k* = 200. Outlier replicates, defined by Euclidean distance, were filtered at a rate of 10%. The consistency of gene expression programs between the data splits was assessed by calculating the Jaccard similarity index of *Z*-score normalized (see Kotlier *et al*.^20^ for details) gene weight coefficients at each value of *k*. *P*-values for all pairwise comparisons between the columns of the component matrices were calculated using the jaccard.test.mca function from the ‘jaccardtest’ R package^49^ followed by Bonferroni multiple testing correction. This was carried out over a range of Jaccard lengths. All significant pairs (*P* < 0.05) of components between data splits were then represented as node pairs in an undirected network graph using the R package *igraph*^50^. Community detection was performed on the underlying graph using edge betweenness centrality, and the number of distinct communities were recorded. These communities represent replicated gene expression programs between data splits, and these were used as the basis of comparison to determine optimal parameters for rank (*k*) and Jaccard length on each dataset. For each Jaccard length the number of discrete communities was plotted against all values of *k*, and a loess regression was performed to facilitate curve smoothing. The inflection point of these concave curves was calculated using the ‘elbow’ R package (https://nicolascasajus.fr/elbow/). Cost values, representing residual values after the inflection point, were used to evaluate the stability of the solution. A linear regression was performed on cost values of each Jaccard length curve, and the slope was recorded. The optimal Jaccard length was established by determining which slope was closest to −1.0, which represents a solution whereby the number of reproducible programs remains approximately constant for all values of *k* greater than ground truth (Supplementary Fig. 1d). Using the established parameters for rank (*k*) and Jaccard length, final reproducible programs for each dataset were calculated by averaging over all nodes (i.e., cNMF gene loading scores (i.e. weights)) per community in the network graph.

### Benchmarking acNMF against other methods

We benchmarked acNMF against 3 different published latent factor models, iNMF, ScVI, and SignatureAnalyzer, using our simulated dataset of 15,000 cells, encompassing 13 cell types plus 1 activity program. We benchmarked all models using the same 2000 highly variable genes. iNMF was run with miniBatch_size = 5000, max.epochs = 5, where we first ran the suggestK function at lambda = 5, thresh = 1e-4 and max.iters = 100 with values of k ranging from 5 to 100 (from 5:60 in intervals of 5, from 70:100 in intervals of 10). We then determined the optimal value of k by LOESS (same approach as in acNMF) on the resulting median KL divergence data. ScVI was run with max_epochs = 200, gene_likelihood = “nb” and latent_distribution = “ln”. k was chosen based on the marginal log-likelihood obtained with the dimensionality of the bottleneck layer ranging from 2 to 100 (from 3:60 in intervals of 1, from 70:100 in intervals of 10), where the encoder and decoder were both structured with 2 layers (note that the marginal log-likelihood fluctuated with increasing dimensionality and we picked the smallest dimensionality with a comparable marginal log-likelihood). SA-GPU was run with K_0 = 100, max_iter = 100000. For ARI (adjusted rand index) calculation, we labeled each cell with the program associated with the highest program expression score (h) and ARI was calculated by ARI function in R library aricode.

### Identifying gene expression programs replicated across different datasets using a network graph

To determine which gene expression programs were replicated across datasets, or between human tumors and preclinical models, all pairwise comparisons between programs from all datasets were performed using an asymmetric Jaccard similarity test, fixing the Jaccard length to that tuned for each dataset above, followed by Bonferroni correction (keeping connections at *FWER* < 0.05). We visually represented connected nodes in a network graph (e.g., Fig. 4a) – the layout of which was established using the force-directed Fruchterman-Reingold algorithm to minimize overlapping of edges. The network structure facilitated the interpretation of these programs, with communities detected by modularity scores calculated from random walks of the graph.

### Web-based platform for annotation, exploration, and cross-dataset integration of acNMF output

For each gene expression program, an interactive web-based report was created, which was used to support annotation and interpretation. To estimate variability between subjective annotators, the GOSH dataset was independently annotated by co-authors P.G. and Y.K, and a consensus annotation was subsequently reached by discussing and merging these annotations, including subjective assignment of confidence levels. The web-based reports were created using the R package *knitr*^51^, which for each program summarized a rich catalogue of biologically relevant information/enrichments, including 282 marker gene lists we curated from the literature. The interactive plots were created using *plotly*^52^. The broader network-based representation comparing gene expression programs across datasets, as well as the tools for organizing the reports and cataloguing/displaying relevant metadata was created using *Shiny*^53^. The complete web platform is accessible at http://pscb.stjude.org.

### RNA *in situ* hybridization (RNA ISH) and HALO analysis

RNA ISH was performed in fresh frozen mouse tissue samples (NB837, NB839, NB853, NB855, NB856, NB864, NB883, NB887) using the RNAscope HiPlex V2 assay Reagent Kit. These data were generated by Advanced Cell Diagnostics (7707 Gateway Blvd., Newark, CA 94560, USA). Briefly, fresh frozen tissue sections were pretreated with 10% NBF and protease prior to hybridization with the target oligo probes (Mm-Tagln, Mm-Acta2, Mm-C3, Mm-Sox10, Mm-Plp1, Hs-MYCN, Mm-Twist1, and Mm-Ddc). Immediately following hybridization, the RNAscope Fluorescent V2 Assay was performed with the preamplifier, amplifier, and RNAscope HiPlex FFPE Reagent to reduce autofluorescence. Fluorophores T1-T4 were then hybridized sequentially, followed by RNAscope DAPI counterstain. After imaging, the first group of multiplexed fluorophores were then cleaved, and the process was repeated for each additional group of fluorophores. Fluorescence images were acquired using a Polaris 40X full-slide digital slide scanner. The resulting multispectral images were then uploaded into HALO Image Analysis platform v3.5.3577.140 (Indica Labs), and the High-Plex FL v2.0 module was used to construct the image analysis algorithms. These included nuclei detection, cytoplasmic segmentation, and FISH detection criteria based on intensity thresholds for each probe in each sample.

### CNV Inference in scRNA-seq data

We used InferCNV^37^ to identify regions of copy number variation from single-cell RNA-seq, which we used as evidence to differentiate between cancer and non-cancerous “normal” cells. InferCNV was run on each patient/sample independently. Reference normal cells were selected from clear immune cells, identified by acNMF (selecting cells expressing an immune program at *h* > 100). In the case of the GOSH dataset, we used the author’s original immune cell annotations, which were essentially identical to the acNMF annotations. In cell lines, we defined the reference normal using immune cells from the Dong dataset. InferCNV’s six-state Hidden Markov Model (i6) predicted CNV regions and states, with posterior probabilities of CNV regions assigned to a given state using a Bayesian network at a value of *P*(Normal) = 0.3 (all other parameters were used at default settings).

### Machine learning classifiers for predicting cancer/normal status of mesenchymal-like cells using inferred copy number profiles

To predict the cancer/normal status of mesenchymal-like cells on a per-cell basis, we tested 3 classification models, a regularized logistic generalized linear model (GLM), a random forest (RF), and a support vector machine (SVM), which all achieved comparable performance. These were implemented in the R packages *glmnet*, *randomForest,* and *SVM* (linear kernel) respectively. We identified cancer cells, endothelial cells, or ambiguous mesenchymal-like cells if an appropriate program was expressed in that cell at *h* > 100. We randomly split clear cancer cells into 75% training set and 25% test set. Cell-specific copy number estimates, which were treated as the model’s input features, were obtained from inferCNV’s predicted regions of copy number variation (inferCNV-features) or were obtained manually, by taking the average standardized TPM expression level in groups of 50 collocated genes along the genome (manual-features). For each sample, a classifier was then trained using sample-specific cancer cells and immune cells (representing normal cells). For each sample, these trained models were then applied to the held-out reference normal endothelial cells, or the held-out ambiguous mesenchymal-like cells, outputting a predicted class of 0 (normal) or 1 (cancer) for each cell. These results were then assembled into a 2 × 2 contingency table, where we applied a Fisher’s exact test to compare the frequency with which the model classified mesenchymal-like cells as cancer cells, against the frequency it misclassified reference normal endothelial cells as cancer cells. Note that for model training/testing in the GEMM data, because the mice were genetically identical, we pooled the immune cells and endothelial cells in the drug-treated and vehicle-treated groups, which increased the size of the reference normal pool of cells and should improve power.

### Differential expression, gene set enrichment analysis, and RNA-velocity analysis of cisplatin treated GEMM

We performed differential expression analysis of our cisplatin treated vs vehicle treated GEMM using the R package DESeq2^54^ by ‘pseudobulking’ cells from each mouse using the aggregate.Matrix function in the R package Matrix.utils. All genes were ranked by logFC between vehicle and cisplatin treated mice, and then were subjected to GSEA^55^ for functional enrichments of the mSigDB Hallmarks gene set, using default parameter settings. Trajectory inference and pseudotime analysis were performed with the R package Monocle3^56^. RNA velocity was assessed by applying the dynamical module of scVelo^56^ (v0.2.5) under default settings.

### Spatial pathology analyses and immunohistochemistry

For H&E staining, the harvested mouse tumor tissue was embedded in Tissue-Tek O.C.T. embedding medium (Sakura Finetek) and flash frozen on a metal block in liquid nitrogen then preserved at –80°C until further processing. Each tissue block was equilibrated to −20°C in a cryostat chamber (CM3050S, Leica BioSciences) for 30 minutes before making 10-µm sections collected onto Superfrost slides. The slides were incubated at 37°C for 1 minute and transferred to a container containing pre-chilled methanol for 30 minutes at −20C in the upright position. After methanol fixing, the excess methanol was wiped out from the bottom of the slides without disrupting the tissue section. After methanol fixation, the slides were immersed into water 10 times gently and incubated in hematoxylin solution for 1 minute followed by immersing slides 10 times into 800 ml water. The slides were then submerged 10 times into Bluing buffer and immersed into water to remove the excess Bluing buffer. Next, slides were dipped into 96% ethanol 10 times followed by incubation with Eosin for 30 seconds. The excess Eosin was removed by immersing slides 10 times into 3 different containers containing 100% ethanol. After that, tissue slides were mounted, and images were acquired using Axioscan slide scanner (Zeiss, Oberkochen, DE) up to a 20X scalable magnification. For our immunohistochemistry analysis of fresh frozen slides used for the RNA ISH experiments serial unstained sections of murine NB-like tumors were fixed in acetone for 20 minutes. Fixed slides were then wet loaded on the Ventana Discovery Ultra autostainer (Roche, Basel, CH) using the Ventana Discovery Ultra antigen retrieval CC1 (catalogue #950-500) for 64 minutes followed by the application of anti-PHOX2B antibody (Abcam, ab183741) at a 1:100 dilution for 60 minutes. OmniMap anti-rabbit HRP (catalogue #760-4311) for 16 minutes, followed by the ChromoMap DAB kit (catalogue #760-159) for 8 minutes, followed by counterstained with hematoxylin, and the post-counterstain Bluing Reagent (catalogue #760-2037) for 8 minutes was used for the detection and visualization of PHOX2B in tissue sections.

Xenograft tissue was fixed in 10% neutral-buffered formalin and processed as paraffin embedded samples. Tissues were sectioned at a 4-µm thickness and mounted onto positively charged glass slides (Superfrost Plus; ThermoFisher Scientific, Waltham, MA). Unstained slides were baked for 60 minutes at 60°C. After deparaffinization, slides underwent immunohistochemical staining for anti-COL1A1 (clone E3E1X, Cell Signaling, #66948) was performed on the Leica BOND-MAX automated stainer (Leica Biosytems, Buffalo Grove, IL) with IHC Protocol F. Heat-induced epitope retrieval was carried out with Bond Epitope Retrieval Solution 2 (ER2) for 20 minutes followed by incubation of the primary antibody at 1:200 for 30 minutes, and visualization with Bond Polymer Refine Detection kit (DS9800). Negative and positive tissue controls included xenografts defined as having either negative or high expression of COL1A1 as determined by sequencing data. Species cross-reactivity was not observed in wild-type B6 murine haired skin, which served as another negative tissue control. An isotype control (Mouse BALB/c IgG2a, BD Pharmingen, 349050) was used to confirm the specificity of immunoreactivity for the monoclonal antibody.

## DECLARATIONS

### Ethics approval and consent to participate

Not applicable.

### Consent for publication

Not applicable.

### Availability of data and materials

Simulated scRNA-seq datasets were obtained through CodeOcean (https://doi.org/10.24433/CO.9044782e-cb96-4733-8a4f-bf42c21399e6) using the Step1_Simulate.ipynb Jupyter Notebook. Default parameters, defined by the implementation of the R package ‘splatter’ on a published scRNA-seq dataset, were used to generate five replicate simulations across three conditions of varying signal:noise levels. The raw readcounts from each of these simulations were subjected to the acNMF approach described below (https://rchapple2.github.io/acNMF/). The gene and cell parameters generated from this code were used to evaluate the recovery of programs following the acNMF analysis. Human datasets were downloaded via the following accession numbers. Dong: GSE137804. Jansky: EGAS00001004388. Kildisiute: EGAD00001008345. Cell line and mouse data deposited in GSE229224 and GSE229226, respectively. PDX deposited in GSE228957.

### Authors’ contributions

RHC and PG developed the methods, analyzed the data and drafted the paper. XL developed the CNV calling method and acNMF benchmarking. SN and JE generated the single-cell data. MIMA, CWL, WCW, YZ performed additional analytical work. YK, MP, HML, ML, SCK, CL, JH, CS, MDJ, TC, WJA performed additional experimental and mouse work. HS performed the pathology work. AGP and MD generate the PDX data and provided analytical support.

## Acknowledgements

Not applicable

