## Supplementary Figures for "An integrated single-cell RNA-seq map of human neuroblastoma tumors and preclinical models uncovers divergent mesenchymal-like gene expression programs"

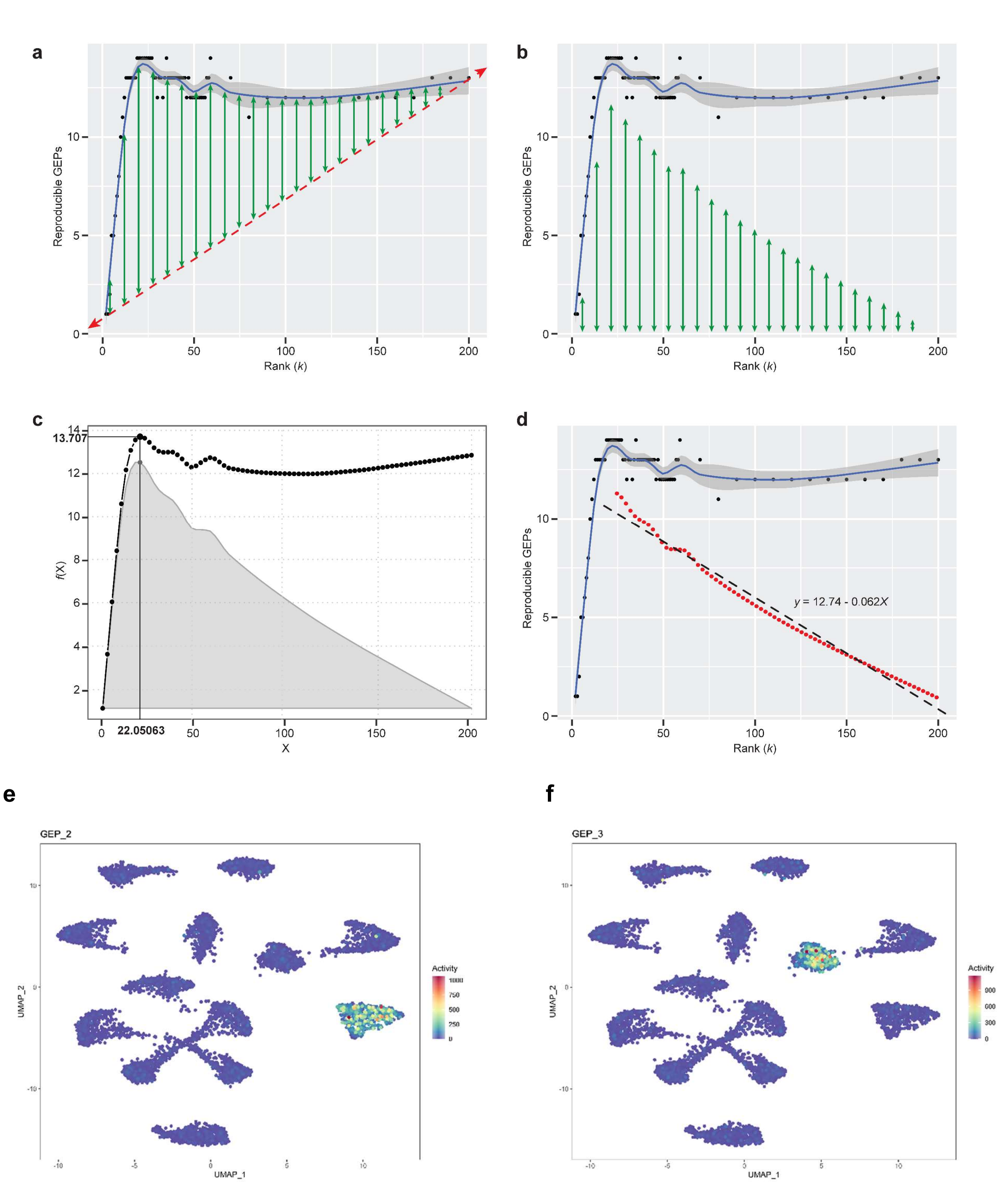
a
b
c
d
e
f
Reproducible GEPs
Rank (k)
f(X)
X
13.707
22.05063
y = 12.74 - 0.062X
GEP_2
GEP_3
UMAP_1
UMAP_2
Activity

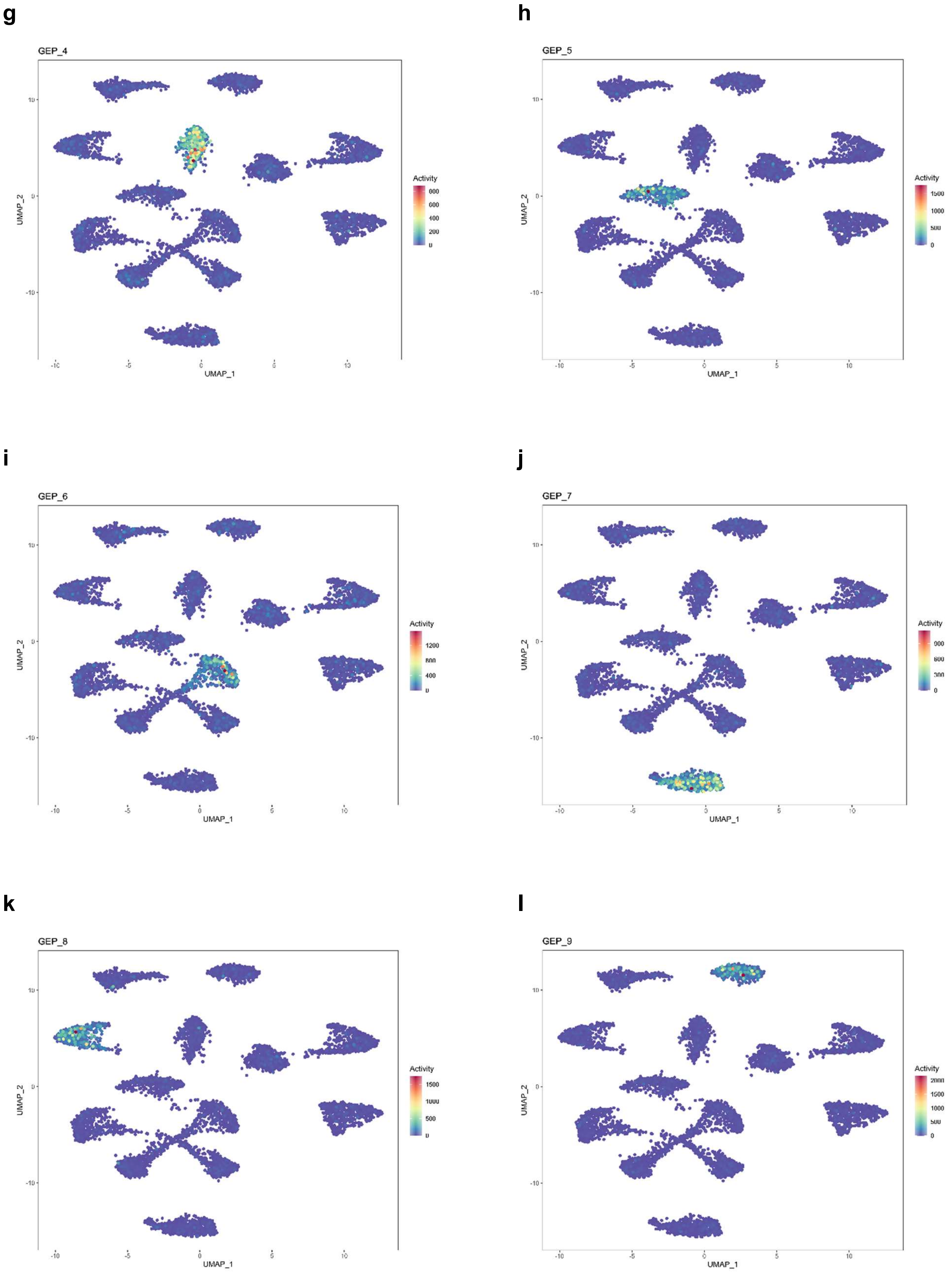

g
GEP_4
h
GEP_5
i
GEP_6
j
GEP_7
k
GEP_8
l
GEP_9
UMAP_1
UMAP_2
Activity

**m**

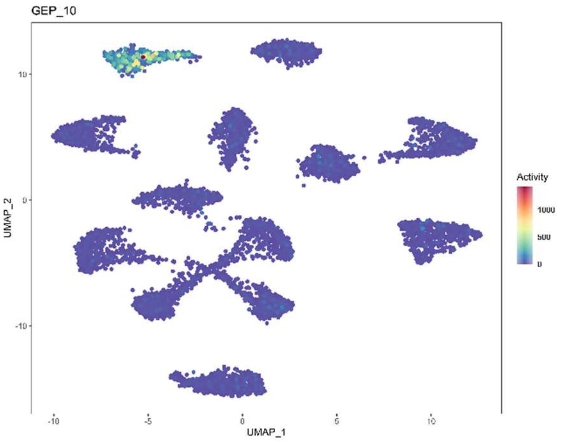
GEP_10
UMAP_2
UMAP_1
Activity
1000
500
0

**n**

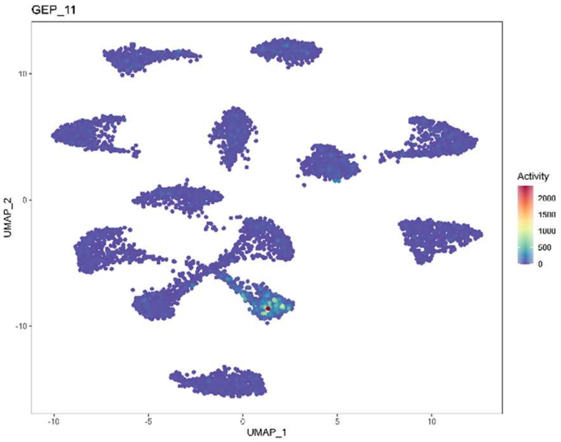
GEP_11
UMAP_2
UMAP_1
Activity
2000
1500
1000
500
0

**o**

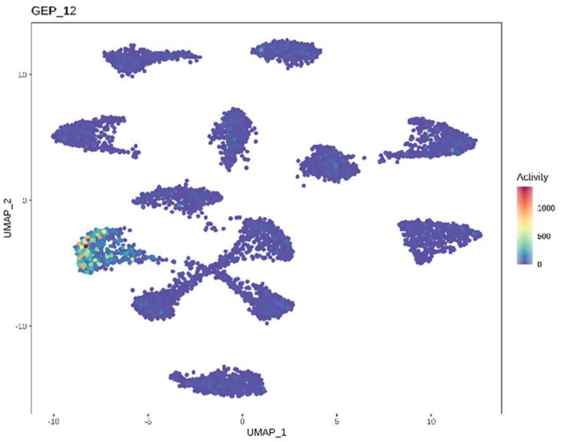
GEP_12
UMAP_2
UMAP_1
Activity
1000
500
0

**p**

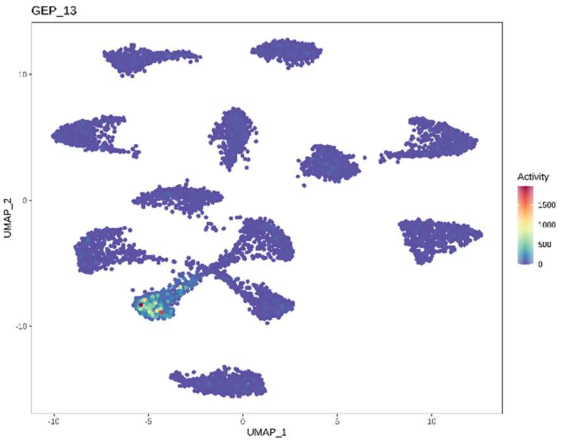
GEP_13
UMAP_2
UMAP_1
Activity
1500
1000
500
0

**q**

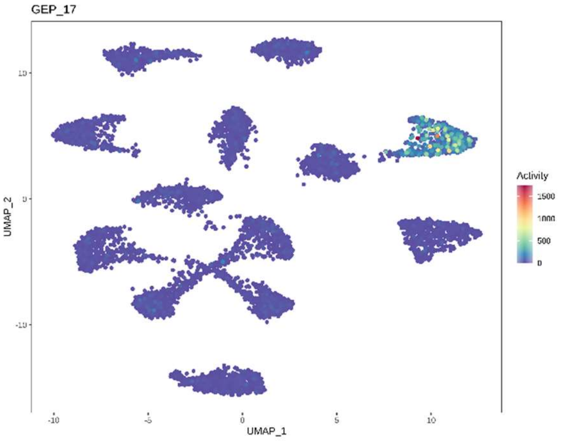
GEP_17
UMAP_2
UMAP_1
Activity
1500
1000
500
0

**r**

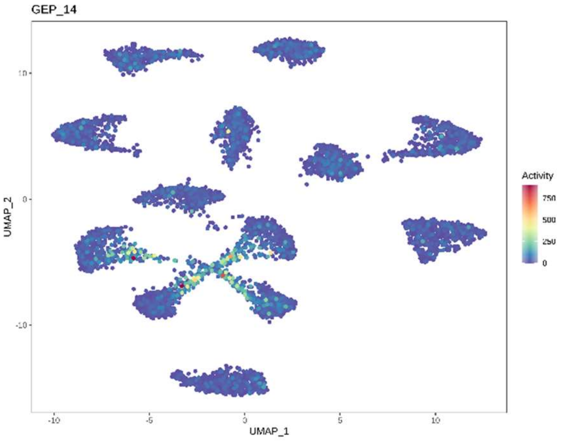
GEP_14
UMAP_2
UMAP_1
Activity
750
500
250
0

**s**

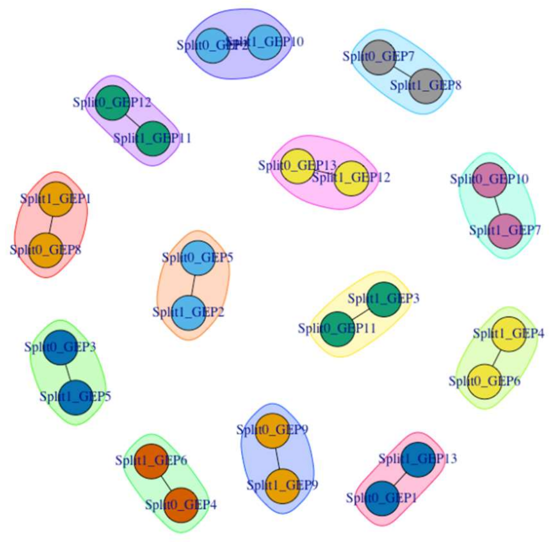


**t**

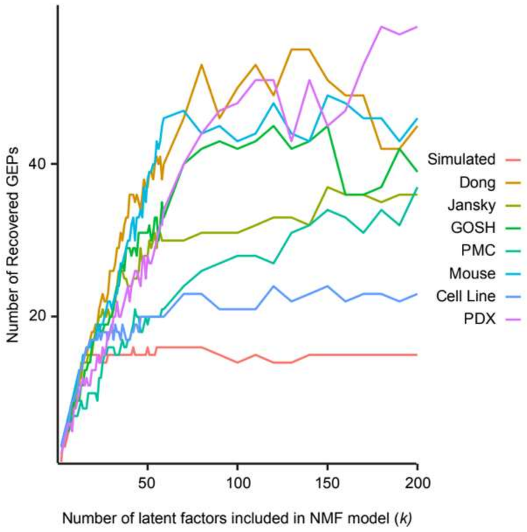


**u**

| | Rank | ARI | K Determination Strategy |
|---|---|---|---|
| acNMF | 14 GEPs @ k = 22 | 0.8879 | |
| iNMF | k = 35 | 0.8294 | suggestK function and inflection point searching algorithm |
| ScVI | k = 36 | 0.8786 | marginal likelihood |
| Signature Analyzer-GPU | K = 21 | 0.7634 | k determined 'automatically' |

**Supplementary Figure 1. acNMF performance on simulated and real data.**

(a-d) Inflection point determination in acNMF curves as implemented by the elbow R package (v0.0.0.9000). cNMF was run across a range of ranks in each independent split of data. Multiple Jaccard lengths were tested as a metric for comparison of the splits, and the resulting network-derived communities (synonymous with reproducible GEPs) were plotted across all ranks. For each Jaccard length, the inflection point of the resulting concave curve is determined by first connecting the minimum and maximum data points (red line) and calculating residuals to the curve (green arrows) (a). After fixing these differences to the x axis, the maximum value of the new profile is chosen (b). This value represents the x value of the inflection point (c), and consequently determines the rank utilized in downstream analyses. The Jaccard length that most reliably stabilizes the solution after the inflection point is represented by fitting a linear regression model of the values on the right side of the inflection point, and choosing the value that minimizes the slope (d).

(e-r) UMAP plots showing all recovered GEPs by acNMF in the simulated single-cell RNA-seq dataset. Dots are colored by the activity score of the recovered gene expression program. All cell identity programs (panels e-q) and the activity program (panel r) were successfully recovered.

(s) Network plot of community detection of simulated data at $k$=14, which results in only 13 reproducible programs (of 14 ground truth programs).

(t) Representative acNMF line plots for each neuroblastoma single-cell RNA-seq dataset highlight the applicability of the method to real data, where in each case the number of replicated gene expression programs reasonably stabilizes beyond a dataset-specific inflection point.

(u) Results of benchmarking acNMF to 3 different published models. A simulated dataset of 15,000 cells of 13 cell types and 1 activity program was used. The adjusted rand index (ARI) was calculated by the ARI function in the R library aricode.

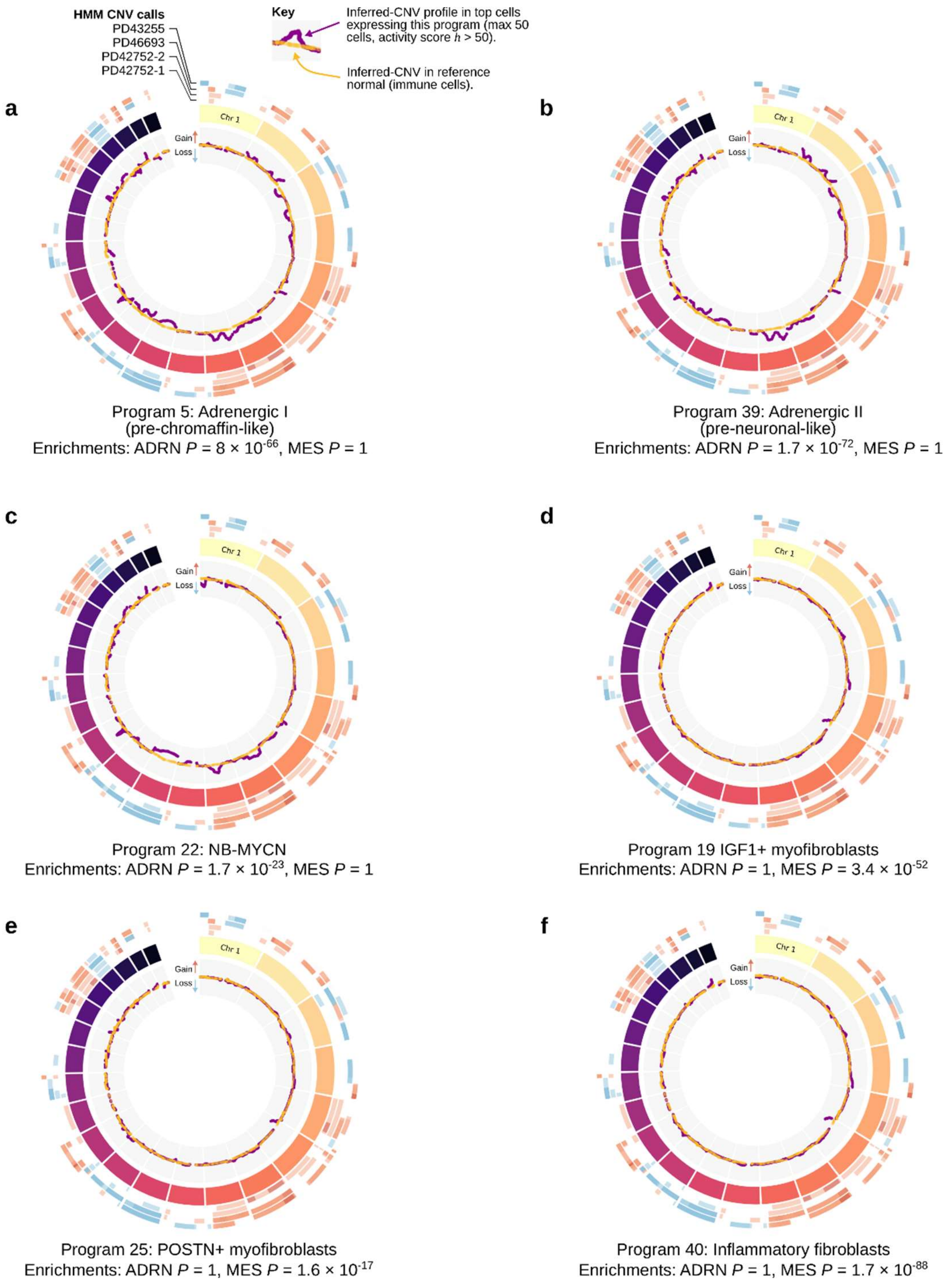


**Supplementary Figure 2. Inferred CNV profiles for gene expression programs enriched for the Van Groningen adrenergic (ADRN) and mesenchymal (MES) signatures in the GOSH dataset.**

(a-f) BioCircos plots representing copy number variation (CNV) profiles inferred from GOSH single-cell RNA-seq data. Each segment on the middle track represents chromosomes 1-22 (ordered clockwise). The outermost track represents Hidden Markov Model (HMM) CNV calls from InferCNV for the high-confidence cancer cells in each of the 4 neuroblastoma samples in GOSH. The innermost track shows the inferred CNV profile in cells highly expressing the program (purple line; calculated from the top 50 cells, or cells with an activity score $h > 50$). The orange line shows the inferred CNV profile for 50 randomly chosen reference normal immune cells. Deviation of the purple line from the orange line is indicative of copy number changes.

Panels (a-c) show the 3 programs with statistical enrichment of the Van Groningen adrenergic (ADRN) signature. Copy number changes are predicted to be abundant in these cells and their expression features are consistent with neuroblastoma cancer cells.

Panels (d-f) show the 3 programs with strongest statistical enrichment of the Van Groningen mesenchymal (MES) signature. Inferred copy number profiles in these cells are largely consistent with the reference normal, and these cells have expression features of fibroblast subpopulations.

NOTE: Our web-based platform allows interactive versions of these plots to be accessed for every gene expression program, across all datasets in this manuscript.

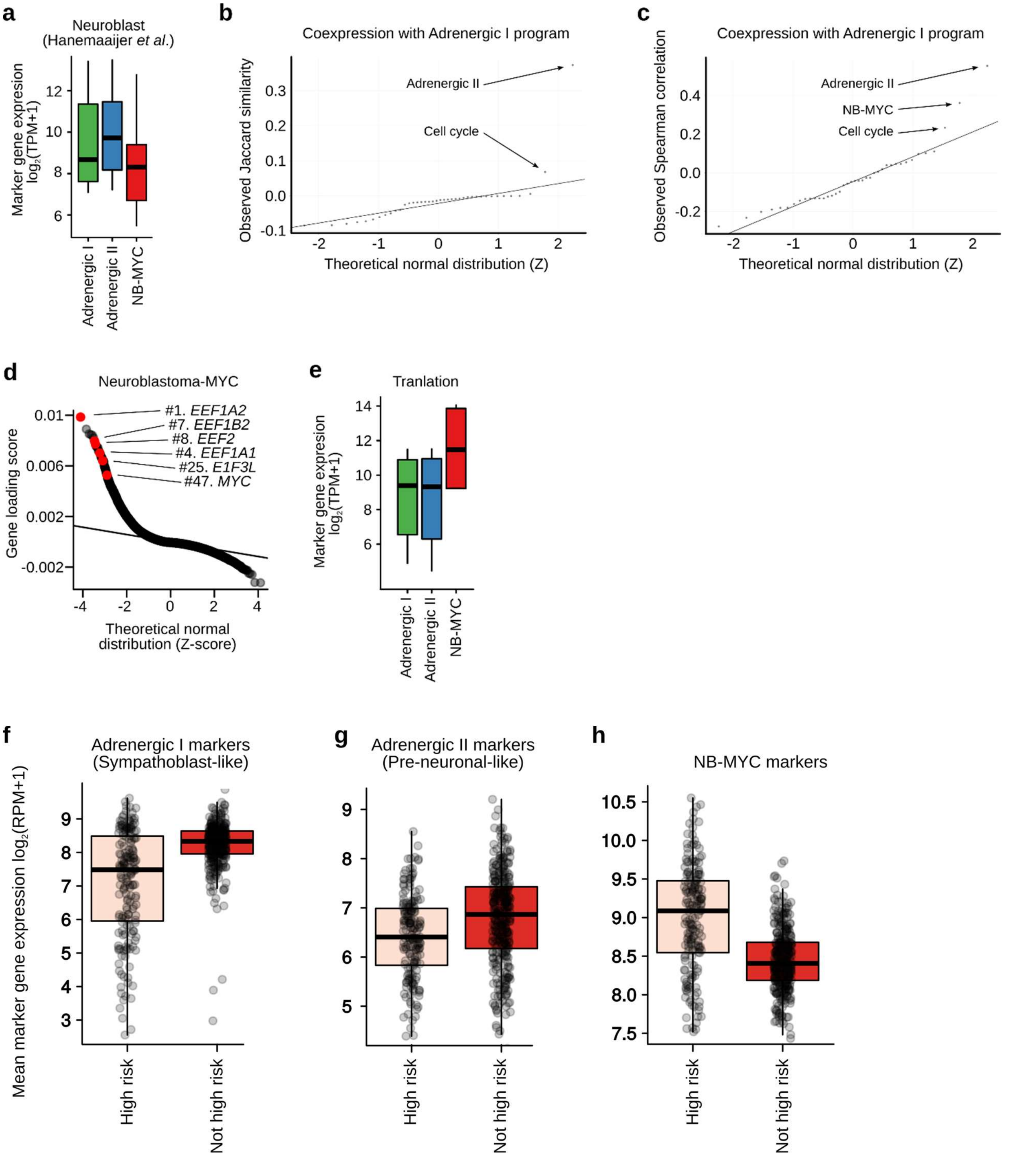


**Supplementary Figure 3. Cells expressing multiple adrenergic and mesenchymal-like gene expression programs are evident in neuroblastoma tumors.**

(a) Boxplots showing the mean gene expression (y axis) of neuroblast marker genes (curated from Hanemaaijer *et al.*) in cells highly expressing ($h > 50$) each of the 3 acNMF recovered adrenergic GEPs (x axis).

(b) QQ-plot summarizing the co-expression of all 40 other GEPs with the Adrenergic I GEP. Co-expression is estimated by an asymmetric Jaccard similarity (y axis), binarizing cells based on whether they express the program at $h > 50$. These values are plotted against a theoretical normal distribution (x axis).

(c) Like (b) but (instead of Jaccard similarity) showing the Spearman's correlation (y axis) of the vectors of the values of $h$ for each program, when compared to the Adrenergic I program.

(d) QQ-plot showing the gene loading scores (y axis) for the "Neuroblastoma-MYC" program. Key genes have been highlighted in red.

(e) Like (a), but for marker genes of "Translation", curated from Dong *et al*.

(f) Mean expression (y axis) of the Adrenergic I (Sympathoblast-like) program using marker genes highlighted in Fig. 3e (*TH*, *DBH*, *CHGA*, *DDC*) in $n = 498$ samples from the SEQ-C bulk RNA-seq dataset (GSE62564).

(g) Like (f) but for the Adrenergic I (pre-neuronal-like) program marker genes highlighted in Fig. 3h (*NEFL*, *NNAT*, *RPRM*, *BASP1*, *NEFM*).

(h) Like (f), but for the NB-MYC program marker genes highlighted in Fig. S3d (*EEF1A2*, *EEF1B2*, *EEF2*, *EEF1A1*, *E1F3L*).

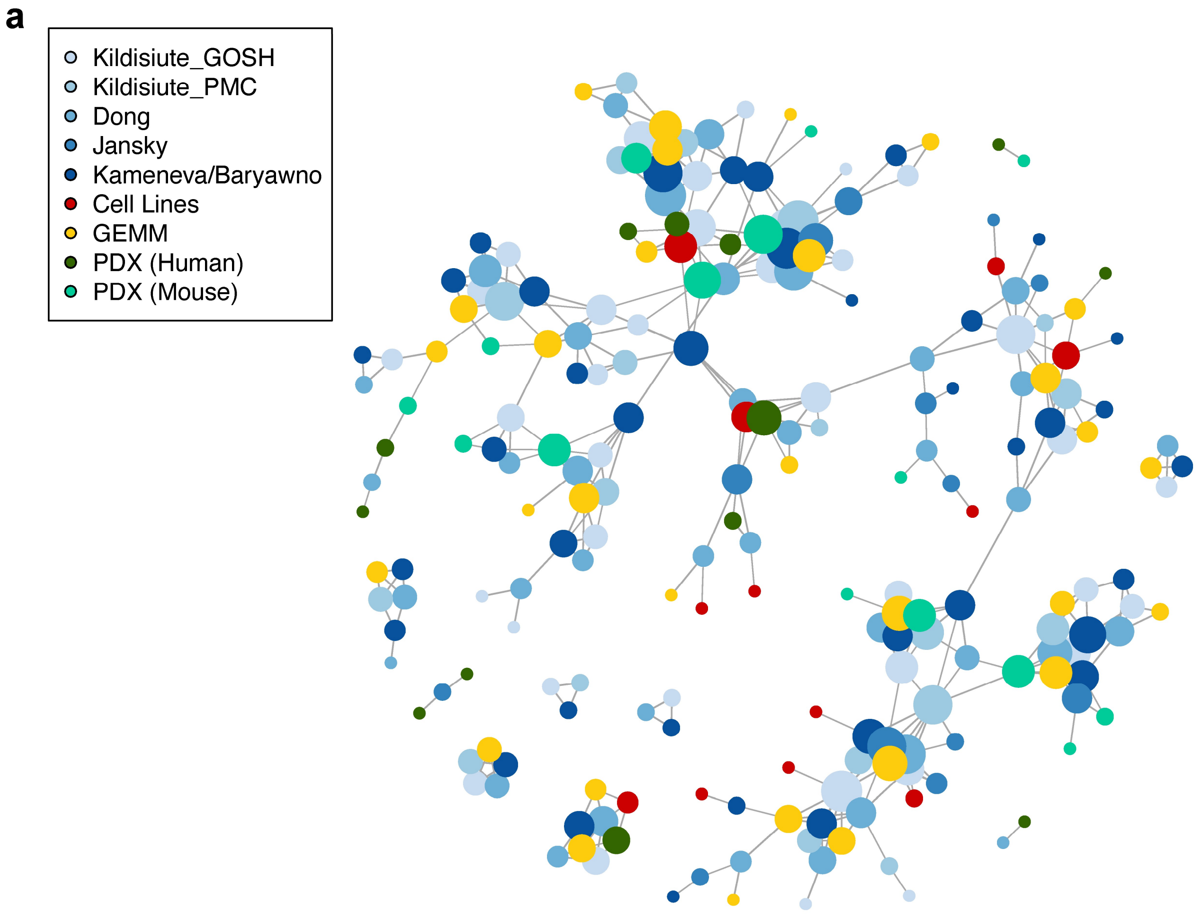


**Supplementary Figure 4. Novel neuroblastoma gene expression programs are reproducible across human neuroblastoma tumor datasets and preclinical models.**

(a) A graph of GEPs identified by acNMF across 8 neuroblastoma single-cell RNA-seq datasets, from humans, PDXs, GEMM, and cell lines. Nodes represent individual GEPs identified through the acNMF method, and edges connect nodes from different datasets if the nodes are statistically matched using a Jaccard similarly test. Node size is represented by the degree of each node. The node connections are identical to Fig. 4a, but here, the nodes have been colored by dataset, rather than by community. Importantly, acNMF identified reproducible GEPs that are not represented in the network because they were not identified in another dataset. This can be partially attributed to patient specific signatures that highlight the ability of acNMF to uncover the distinct heterogeneity of neuroblastoma. Of note, an interactive version of this graph-based representation of these datasets can be accessed in our web-based platform. Each node is clickable and opens a detailed report and analysis of each gene expression program.  Furthermore, the unique GEPs not represented in the network can be found and explored using this browser.

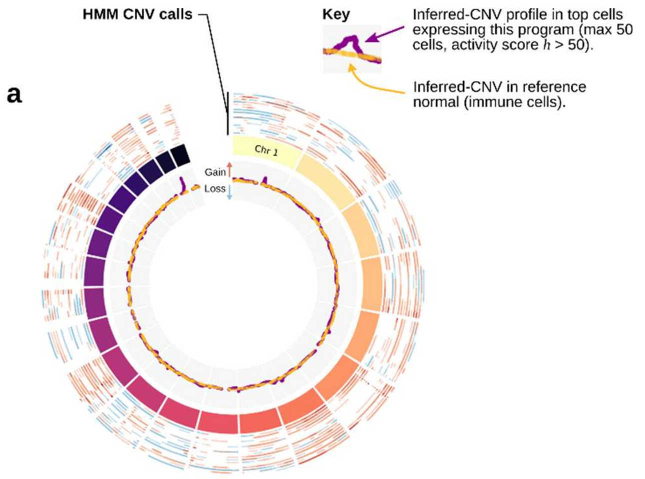


Dong Program 27: Schwann cells
Enrichments: Decartes Schwann cells $P = 1.03 \times 10^{-9}$
ADRN $P = 1$, MES $P = 5.4 \times 10^{-43}$

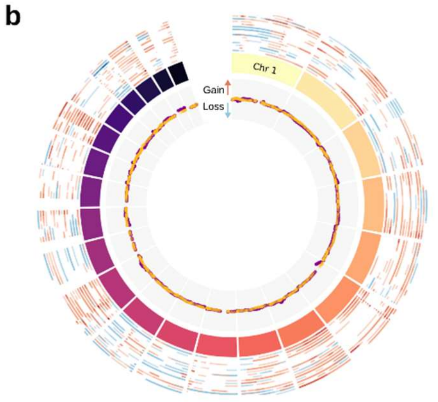


Dong Program 6: Inflammatory Fibroblasts
Enrichments: Lavie inflammatory fibroblasts $P = 2.61 \times 10^{-11}$
ADRN $P = 1$, MES $P = 4.2 \times 10^{-68}$

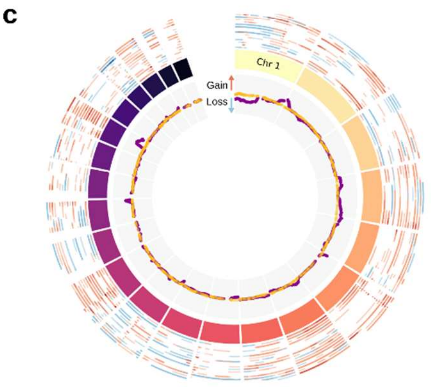


Dong Program 19: Adrenergic I (preneuronal)
ADRN $P = 8.4 \times 10^{-18}$, MES $P = 1$

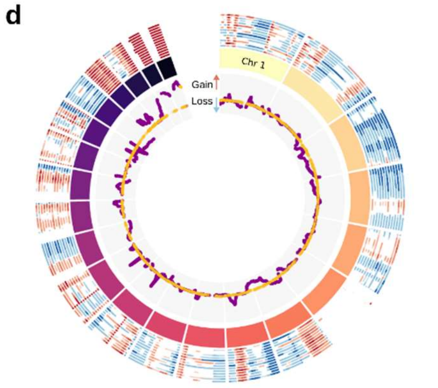


Cell lines Program 14: Adrenergic (BE2M17 / SKNSH)
ADRN $P = 5.2 \times 10^{-34}$, MES $P = 1$

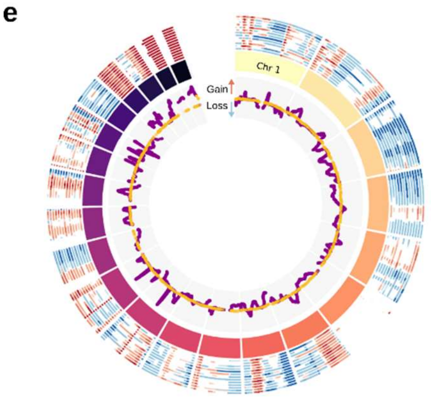


Cell lines Program 7: MES (primarily GIMEN)
ADRN $P = 1$, MES $P = 2.06 \times 10^{-41}$

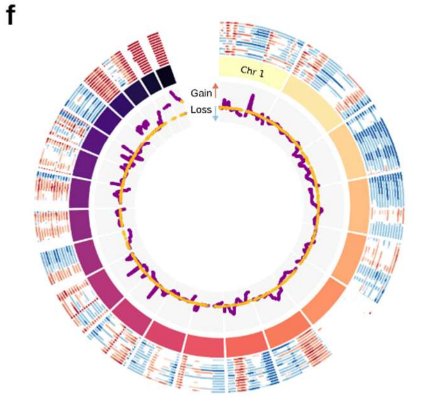


Cell lines Program 8: MES (primarily SKNAS)
ADRN $P = 1$, MES $P = 2.05 \times 10^{-28}$

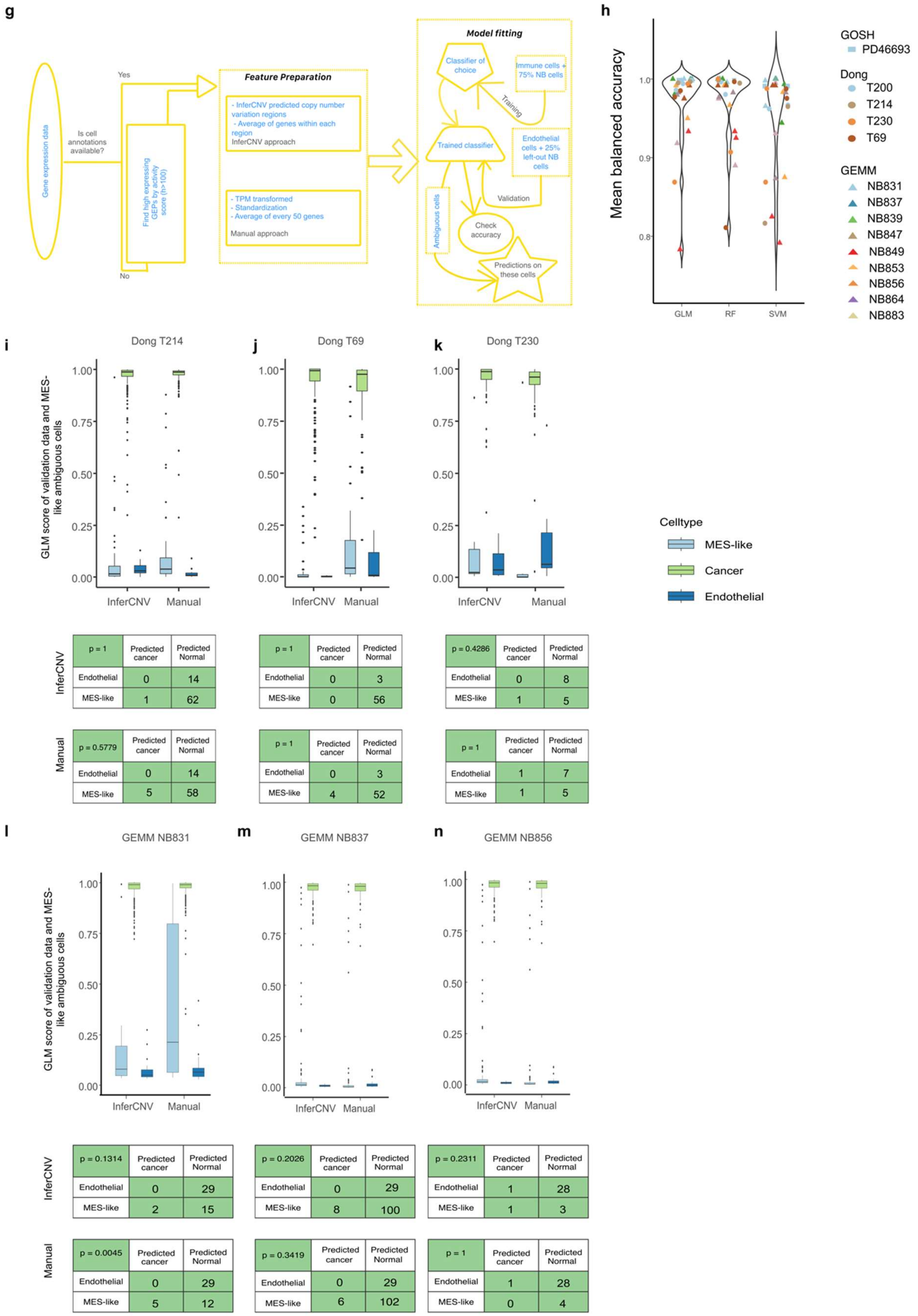
g
Gene expression data
Is cell annotations available?
Yes
No
Find high expressing GEPs by activity score (h>100)
Feature Preparation
- InferCNV predicted copy number variation regions
- Average of genes within each region
InferCNV approach
- TPM transformed
- Standardization
- Average of every 50 genes
Manual approach
Model fitting
Classifier of choice
Immune cells + 75% NB cells
Training
Trained classifier
Endothelial cells + 25% left-out NB cells
Validation
Ambiguous cells
Check accuracy
Predictions on these cells
h
Mean balanced accuracy
GLM
RF
SVM
GOSH
PD46693
Dong
T200
T214
T230
T69
GEMM
NB831
NB837
NB839
NB847
NB849
NB853
NB856
NB864
NB883
i Dong T214
j Dong T69
k Dong T230
GLM score of validation data and MES-like ambiguous cells
InferCNV
Manual
Celltype
MES-like
Cancer
Endothelial
InferCNV, Dong T214: p = 1 | Predicted cancer | Predicted Normal; Endothelial 0 14; MES-like 1 62
InferCNV, Dong T69: p = 1 | Predicted cancer | Predicted Normal; Endothelial 0 3; MES-like 0 56
InferCNV, Dong T230: p = 0.4286 | Predicted cancer | Predicted Normal; Endothelial 0 8; MES-like 1 5
Manual, Dong T214: p = 0.5779 | Predicted cancer | Predicted Normal; Endothelial 0 14; MES-like 5 58
Manual, Dong T69: p = 1 | Predicted cancer | Predicted Normal; Endothelial 0 3; MES-like 4 52
Manual, Dong T230: p = 1 | Predicted cancer | Predicted Normal; Endothelial 1 7; MES-like 1 5
l GEMM NB831
m GEMM NB837
n GEMM NB856
InferCNV, GEMM NB831: p = 0.1314 | Predicted cancer | Predicted Normal; Endothelial 0 29; MES-like 2 15
InferCNV, GEMM NB837: p = 0.2026 | Predicted cancer | Predicted Normal; Endothelial 0 29; MES-like 8 100
InferCNV, GEMM NB856: p = 0.2311 | Predicted cancer | Predicted Normal; Endothelial 1 28; MES-like 1 3
Manual, GEMM NB831: p = 0.0045 | Predicted cancer | Predicted Normal; Endothelial 0 29; MES-like 5 12
Manual, GEMM NB837: p = 0.3419 | Predicted cancer | Predicted Normal; Endothelial 0 29; MES-like 6 102
Manual, GEMM NB856: p = 1 | Predicted cancer | Predicted Normal; Endothelial 1 28; MES-like 0 4

**Supplementary Figure 5. Mesenchymal-like gene expression programs *in vivo*.**

(a-f) BioCircos plots representing copy number variation (CNV) profiles inferred from single-cell RNA-seq data from either the Dong or Cell Line datasets. Each segment on the middle track represents chromosomes 1-22 (ordered clockwise). The outermost track represents Hidden Markov Model (HMM) CNV calls from InferCNV for the high-confidence cancer cells in each of the neuroblastoma samples in GOSH. The innermost track shows the inferred CNV profile in cells highly expressing the program (purple line; calculated from the top 50 cells, or cells with an activity score $h > 50$). The orange line shows the inferred CNV profile for 50 randomly chosen reference normal immune cells.

(a-b) The (heavily mesenchymal-like) Schwann and Fibroblast programs from the Dong dataset, where the CNV profiles are close to the reference normal. Panel c shows one of the Dong adrenergic programs, which deviates from the reference normal.

(d-f) Similar plots from the Cell Line dataset, where both mesenchymal and adrenergic programs deviate heavily from the reference normal.

(g) Flowchart describing the computational procedure for predicting cancer/normal status of cells identified in neuroblastoma tumors expressing a mesenchymal-like program.

(h) A comparison of the balanced accuracy (y axis) achieved predicting cancer/normal status of known cancer or immune cells for the 3 different initial models (x axis) across 15 different tumor samples (points; shown for both “inferCNV” and “manual” features) from either the GOSH, Dong or GEMM datasets. 25% of cancer cells were held out and endothelial cells were treated as high confidence normal calls. GLM, generalized linear model; RF, random forest; SVM, support vector machine. This is similar to Fig. 5(a), but here each tumor is highlighted by color.

(i-o) Boxplots showing the predicted probability of each cell being a cancer cell from the GLM model (y axis) for the ambiguous-mesenchymal cells (light blue), for high confidence cancer cells held out from the initial model fitting (green) and for normal endothelial cells held out from the initial model fitting (dark blue). The predictions are shown for models using features defined based on InferCNV (left 3 boxes) or from manual bins of 50 adjacent genes (right 3 boxes). The panels show the results for samples that contained sufficient numbers of the relevant cell types for this analysis. The contingency tables (right panels) estimate whether the number of mesenchymal-like cells classified as cancer cells differs statistically from the misclassification rate of endothelial cells, calculated by a Fisher's exact test.

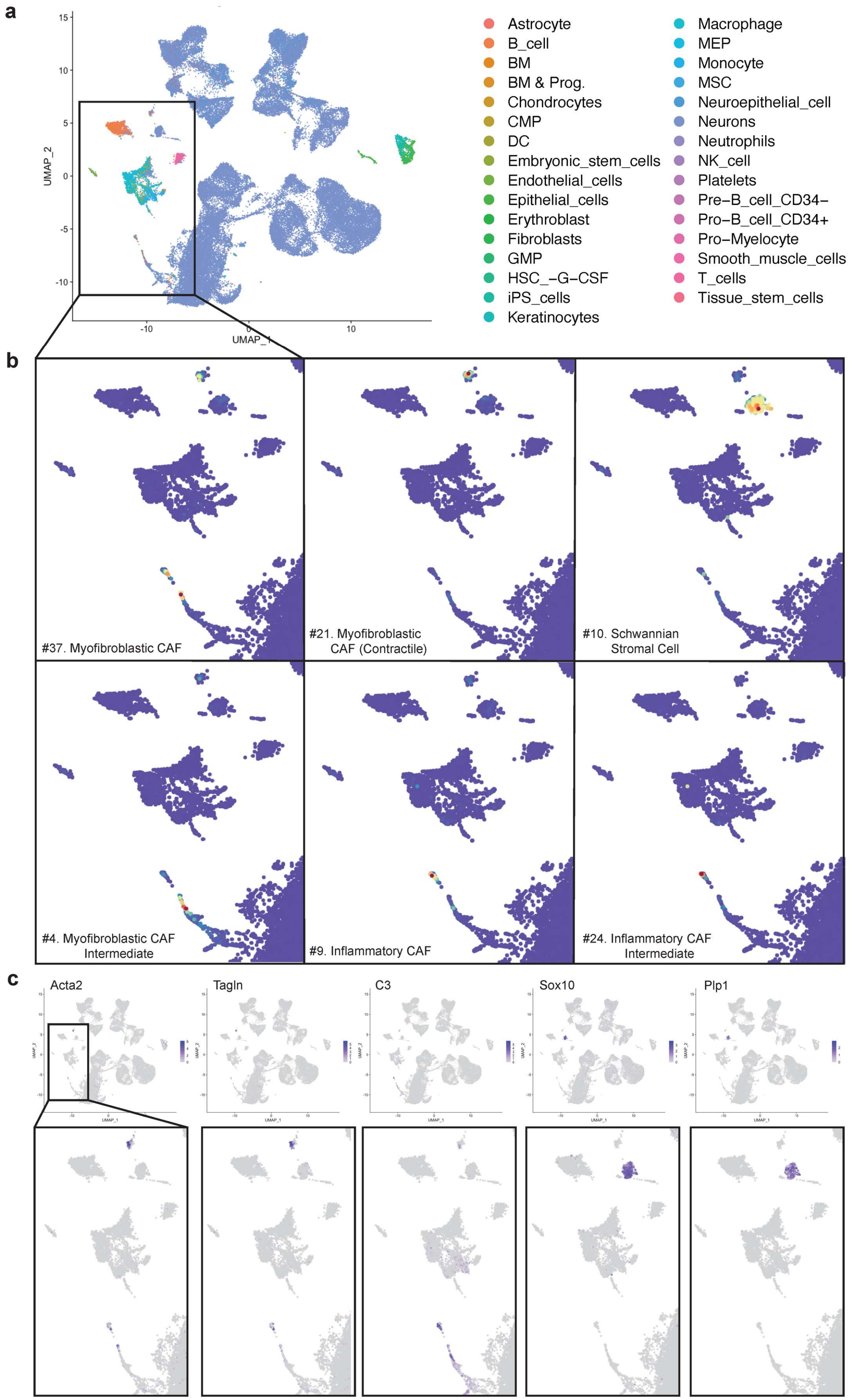
a
UMAP_2
UMAP_1
Astrocyte
B_cell
BM
BM & Prog.
Chondrocytes
CMP
DC
Embryonic_stem_cells
Endothelial_cells
Epithelial_cells
Erythroblast
Fibroblasts
GMP
HSC_-G-CSF
iPS_cells
Keratinocytes
Macrophage
MEP
Monocyte
MSC
Neuroepithelial_cell
Neurons
Neutrophils
NK_cell
Platelets
Pre-B_cell_CD34-
Pro-B_cell_CD34+
Pro-Myelocyte
Smooth_muscle_cells
T_cells
Tissue_stem_cells
b
#37. Myofibroblastic CAF
#21. Myofibroblastic CAF (Contractile)
#10. Schwannian Stromal Cell
#4. Myofibroblastic CAF Intermediate
#9. Inflammatory CAF
#24. Inflammatory CAF Intermediate
c
Acta2
Tagln
C3
Sox10
Plp1

**Supplementary Figure 6. Mesenchymal program identification in mouse NB tumors.**

(a) 2D representation (UMAP) of single cell RNA-seq data from 11 NB tumors in Dbh-iCre; LSL-MYCN mice. Cells are colored by cell type annotation as identified by comparing their expression values to the Human Primary Cell Atlas with SingleR. The highlighted region (bottom left) contains non-malignant cell types from these tumors.

(b) Visualization of acNMF mesenchymal GEPs. Usage scores for GEPs enriched for the Van Groningen mesenchymal signature (from Figure 5d) are plotted in the zoomed region of the UMAP in a). Despite neuroblastoma mesenchymal marker gene expression, these signatures are expressed in other normal cell types. However, in some instances cells can express multiple mesenchymal GEPs simultaneously.

(c) Expression values for individual genes to be used as RNA-ISH probes. Candidate marker genes for cells expressing mesenchymal features were chosen by first investigating GEP loading (weights) scores followed by inspection of read counts.

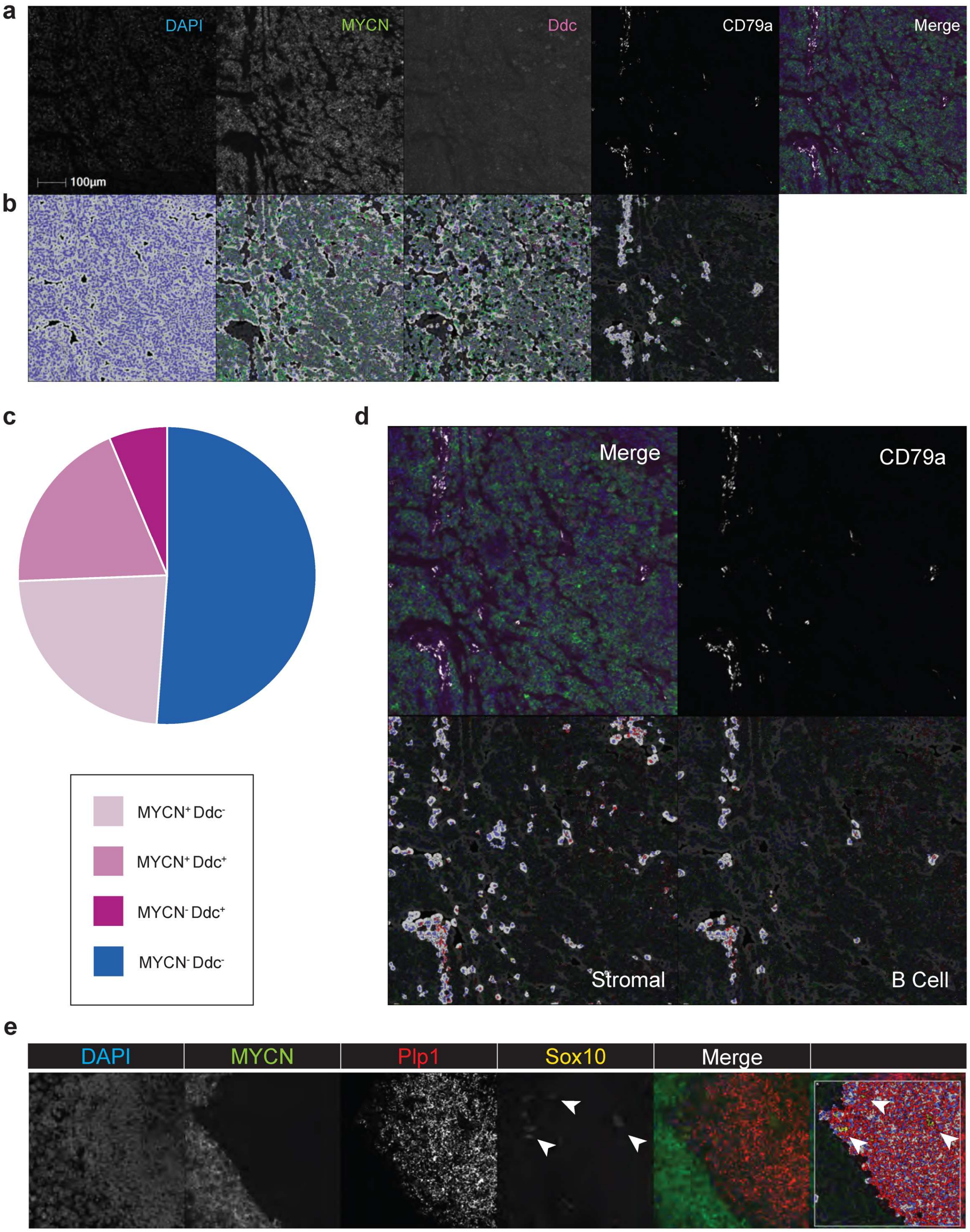

a
DAPI
MYCN
Ddc
CD79a
Merge
100µm
b
c
MYCN+ Ddc-
MYCN+ Ddc+
MYCN- Ddc+
MYCN- Ddc-
d
Merge
CD79a
Stromal
B Cell
e
DAPI
MYCN
Plp1
Sox10
Merge

**Supplementary Figure 7. Analysis of RNA-ISH multispectral data with HALO**

(a-b) Identifying expression status of FISH probes. a) Panels of individual control probes. DAPI counterstaining (leftmost panel in a) facilitated nuclei detection and cytoplasmic segmentation (leftmost panel in b) in HALO analysis software. Analysis of individual gene expression was then performed using fluorescence intensity values (set for each gene in each sample). The fluorescence images in each individual channel are shown in a) and are accompanied by ISH detection by HALO below in b). The ISH detection represents cells that stained positively for that probe. The control probes included two neuroblastoma markers (*MYCN* and *Ddc*) as well as the B cell marker *CD79a*.

(c) Quantification of cells expressing control probes. For each cell in each sample, the status of control probes was assessed. The pie chart exhibits the proportions of cells co-expressing NB markers in *CD79a*-positive cells. Those proportions colored in blue are *CD79a*-positive cells that do not co-express NB markers.

(d) Visual inspection of raw fluorescent images in individual channels and registered images indicated that the colocalization of *CD79a* with NB markers was due to a technical limitation of the analysis software. More specifically, the lack of clear cell outlines results in improper cytoplasmic rendering for any associated nucleus. As such, the NB marker detection in CD79a-expressing cells occurred at tumor/stroma interfaces and are likely false positives. Nevertheless, the majority of *CD79a*-positive cells were negative for both NB markers (shown in blue in Figure c.) Accordingly, the exclusion of both NB markers established the criteria from which the stromal phenotype was derived (bottom left). The pattern of stromal cells matches the regions demarcated by the absence of NB markers (top left). Moreover, by restricting *CD79a* detection in stromal cells, the pattern of B cells can be reliably recovered (bottom right).

(e.) The analysis settings defined in (a-d) were applied to known Schwannian cell markers including *Plp1* and *Sox10*. We observed co-localization of both canonical Schwannian markers in stromal regions (*MYCN* and *Ddc* negative), which can be observed from raw intensity (Merge) or from the HALO rendering (rightmost panel).

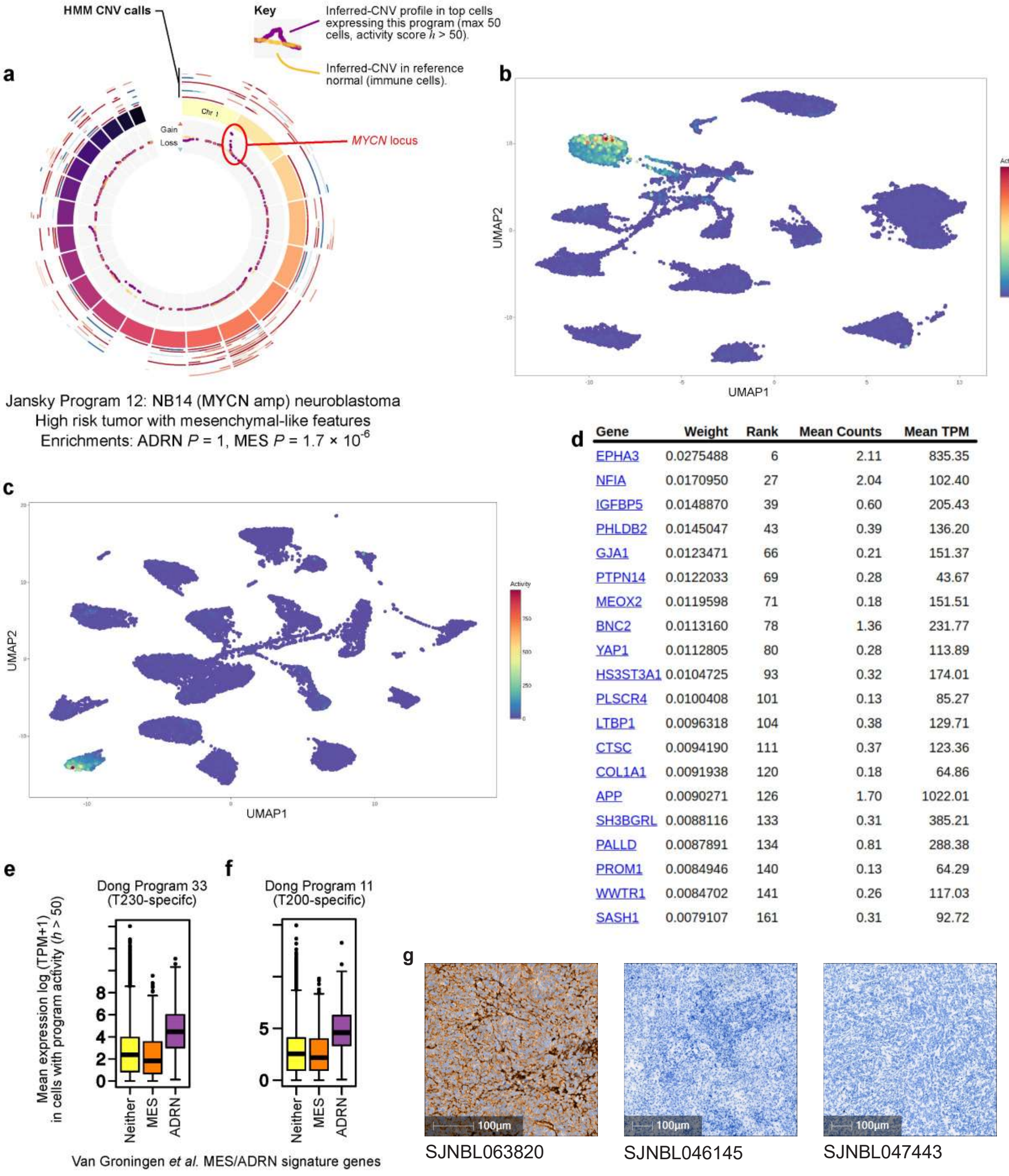


| Gene | Weight | Rank | Mean Counts | Mean TPM |
|---|---|---|---|---|
| EPHA3 | 0.0275488 | 6 | 2.11 | 835.35 |
| NFIA | 0.0170950 | 27 | 2.04 | 102.40 |
| IGFBP5 | 0.0148870 | 39 | 0.60 | 205.43 |
| PHLDB2 | 0.0145047 | 43 | 0.39 | 136.20 |
| GJA1 | 0.0123471 | 66 | 0.21 | 151.37 |
| PTPN14 | 0.0122033 | 69 | 0.28 | 43.67 |
| MEOX2 | 0.0119598 | 71 | 0.18 | 151.51 |
| BNC2 | 0.0113160 | 78 | 1.36 | 231.77 |
| YAP1 | 0.0112805 | 80 | 0.28 | 113.89 |
| HS3ST3A1 | 0.0104725 | 93 | 0.32 | 174.01 |
| PLSCR4 | 0.0100408 | 101 | 0.13 | 85.27 |
| LTBP1 | 0.0096318 | 104 | 0.38 | 129.71 |
| CTSC | 0.0094190 | 111 | 0.37 | 123.36 |
| COL1A1 | 0.0091938 | 120 | 0.18 | 64.86 |
| APP | 0.0090271 | 126 | 1.70 | 1022.01 |
| SH3BGRL | 0.0088116 | 133 | 0.31 | 385.21 |
| PALLD | 0.0087891 | 134 | 0.81 | 288.38 |
| PROM1 | 0.0084946 | 140 | 0.13 | 64.29 |
| WWTR1 | 0.0084702 | 141 | 0.26 | 117.03 |
| SASH1 | 0.0079107 | 161 | 0.31 | 92.72 |

**Supplementary Figure 8. Rare mesenchymal-like signatures in neuroblastoma cells in human and PDX models.**

(a) BioCircos plot (similar to panels in Supplementary Figure 5 (a-f)), but for Program 12 in the Jansky dataset, which is exclusive to the NB14 sample and weakly expresses mesenchymal-like features (while remaining predominantly adrenergic).

(b) UMAP plot showing the activity of Program 12 in the Jansky dataset.

(c) UMAP plot showing the activity of Program 19 in the PDX dataset, (specific to SJNBL063820; which also weakly expresses mesenchymal-like features).

(d) Table showing the gene weights (loadings) for the top-ranked Van Groningen mesenchymal signature genes on Program 19 from the PDX dataset (i.e. these correspond to the highly differentially expressed genes in cells highly expressing Program 19).

(e) Boxplots showing the mean expression (y axis) of the genes from the classical Van Groningen mesenchymal (MES; orange box) or adrenergic (ADRM; purple box) signature, in cells that are highly expressing ($h > 50$) the acNMF-identified *TWISTT1* expressing Program 33 in the Dong dataset, which is expressed in cancer cells from the T230 sample. The yellow box shows the mean expression of all other genes in these cells.

(f) Like (t) but for Program 11, which is expressed in cancer cells of the T200 sample in Dong.

NOTE: Our web-based platform allows interactive versions of these plots, along with a very detailed report, to be accessed for every gene expression program, across all datasets in this manuscript.

(g) Full sized immunohistochemical staining images for the mesenchymal-marker gene COL1A1 in the three PDX samples from Figure 5j.

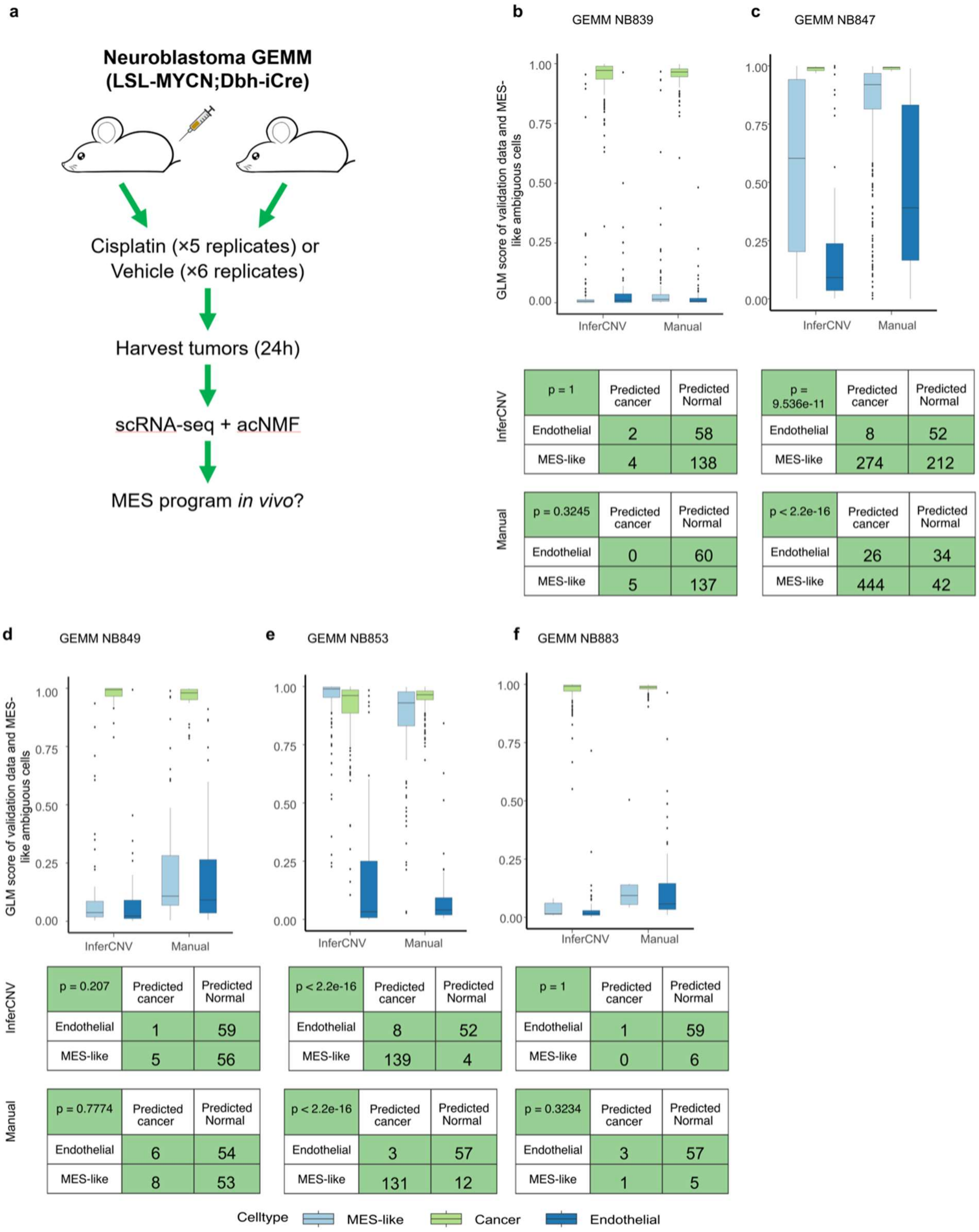


**b, GEMM NB839 — InferCNV**

| p = 1 | Predicted cancer | Predicted Normal |
|---|---|---|
| Endothelial | 2 | 58 |
| MES-like | 4 | 138 |

**b, GEMM NB839 — Manual**

| p = 0.3245 | Predicted cancer | Predicted Normal |
|---|---|---|
| Endothelial | 0 | 60 |
| MES-like | 5 | 137 |

**c, GEMM NB847 — InferCNV**

| p = 9.536e-11 | Predicted cancer | Predicted Normal |
|---|---|---|
| Endothelial | 8 | 52 |
| MES-like | 274 | 212 |

**c, GEMM NB847 — Manual**

| p < 2.2e-16 | Predicted cancer | Predicted Normal |
|---|---|---|
| Endothelial | 26 | 34 |
| MES-like | 444 | 42 |

**d, GEMM NB849 — InferCNV**

| p = 0.207 | Predicted cancer | Predicted Normal |
|---|---|---|
| Endothelial | 1 | 59 |
| MES-like | 5 | 56 |

**d, GEMM NB849 — Manual**

| p = 0.7774 | Predicted cancer | Predicted Normal |
|---|---|---|
| Endothelial | 6 | 54 |
| MES-like | 8 | 53 |

**e, GEMM NB853 — InferCNV**

| p < 2.2e-16 | Predicted cancer | Predicted Normal |
|---|---|---|
| Endothelial | 8 | 52 |
| MES-like | 139 | 4 |

**e, GEMM NB853 — Manual**

| p < 2.2e-16 | Predicted cancer | Predicted Normal |
|---|---|---|
| Endothelial | 3 | 57 |
| MES-like | 131 | 12 |

**f, GEMM NB883 — InferCNV**

| p = 1 | Predicted cancer | Predicted Normal |
|---|---|---|
| Endothelial | 1 | 59 |
| MES-like | 0 | 6 |

**f, GEMM NB883 — Manual**

| p = 0.3234 | Predicted cancer | Predicted Normal |
|---|---|---|
| Endothelial | 3 | 57 |
| MES-like | 1 | 5 |

**Supplementary Figure 9. Mesenchymal-like gene expression programs in drug treated tumors.**

(a) Schematic representation of our experimental design, where single-cell RNA-seq profiles in drug-treated tumors were compared to vehicle-treated tumors.

(b-f) Boxplots showing the predicted probability of each cell being a cancer cell from the GLM model (y axis) for the ambiguous-mesenchymal cells (light blue), for high confidence cancer cells held out from the initial model fitting (green) and for normal endothelial cells held out from the initial model fitting (dark blue). The predictions are shown for models using features defined based on InferCNV (left 3 boxes) or from manual bins of 50 adjacent genes (right 3 boxes). The panels show the results for mesenchymal-like programs in the 6 drug-treated mice. The contingency tables (lower panels) estimate whether the number of mesenchymal-like cells classified as cancer cells differs statistically from the misclassification rate of endothelial cells, calculated by Fisher's exact test.

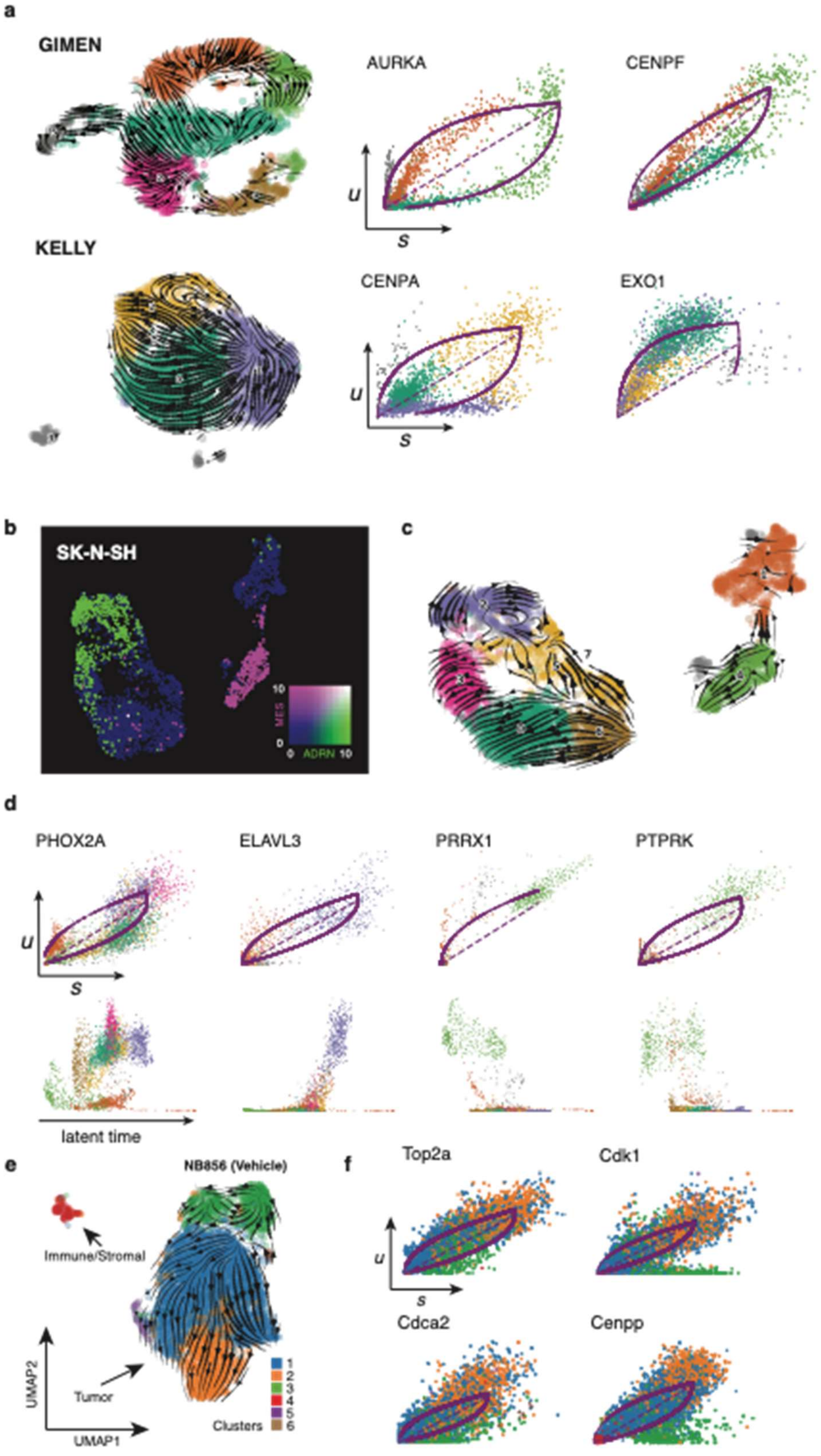

a
GIMEN
AURKA
CENPF
u
s
KELLY
CENPA
EXO1
b
SK-N-SH
MES
ADRN
c
d
PHOX2A
ELAVL3
PRRX1
PTPRK
latent time
e
NB856 (Vehicle)
Immune/Stromal
Tumor
UMAP2
UMAP1
Clusters
1
2
3
4
5
6
f
Top2a
Cdk1
Cdca2
Cenpp

**Supplementary Figure 10. Examining transcriptional dynamics in cell lines with RNA velocity.**

(a) RNA velocity was estimated in both mesenchymal and adrenergic neuroblastoma cell lines. While examples of both mesenchymal (GIMEN, upper panel) and adrenergic (KELLY, lower panel) cell lines exhibit large changes in gene expression, the main drivers of these RNA velocity vectors are cell cycle regulators and proliferative signals.

(b) Simultaneous expression of adrenergic (green) and mesenchymal (magenta) signatures are observed in the SK-N-SH neuroblastoma cell lines as described previously (Thirant et. al., 2023, Boeva et al. 2017)

(c) RNA velocity vectors in the admixed SK-N-SH neuroblastoma cell line indicates two modes of regulation in two phenotypically divergent groups of cells.

(d) Transcriptional dynamics underpinning the characteristics of the vectors found in (c) reveal genes indicative of adrenergic and mesenchymal cell states, respectively. Both the RNA velocity plots (upper panel) and the pseudotemporal plots (lower panel) indicate that these genes involved in cellular identity are undergoing significant turnover in distinct groups of cells.

(e) RNA velocity on representative example of our vehicle-treated GEMM.

(f) RNA velocity plots of high likelihood genes in the vechicle treated mice. Similar to NB cell lines at homeostasis, most of the transcriptional dynamics are attributable to cell cycle regulators.

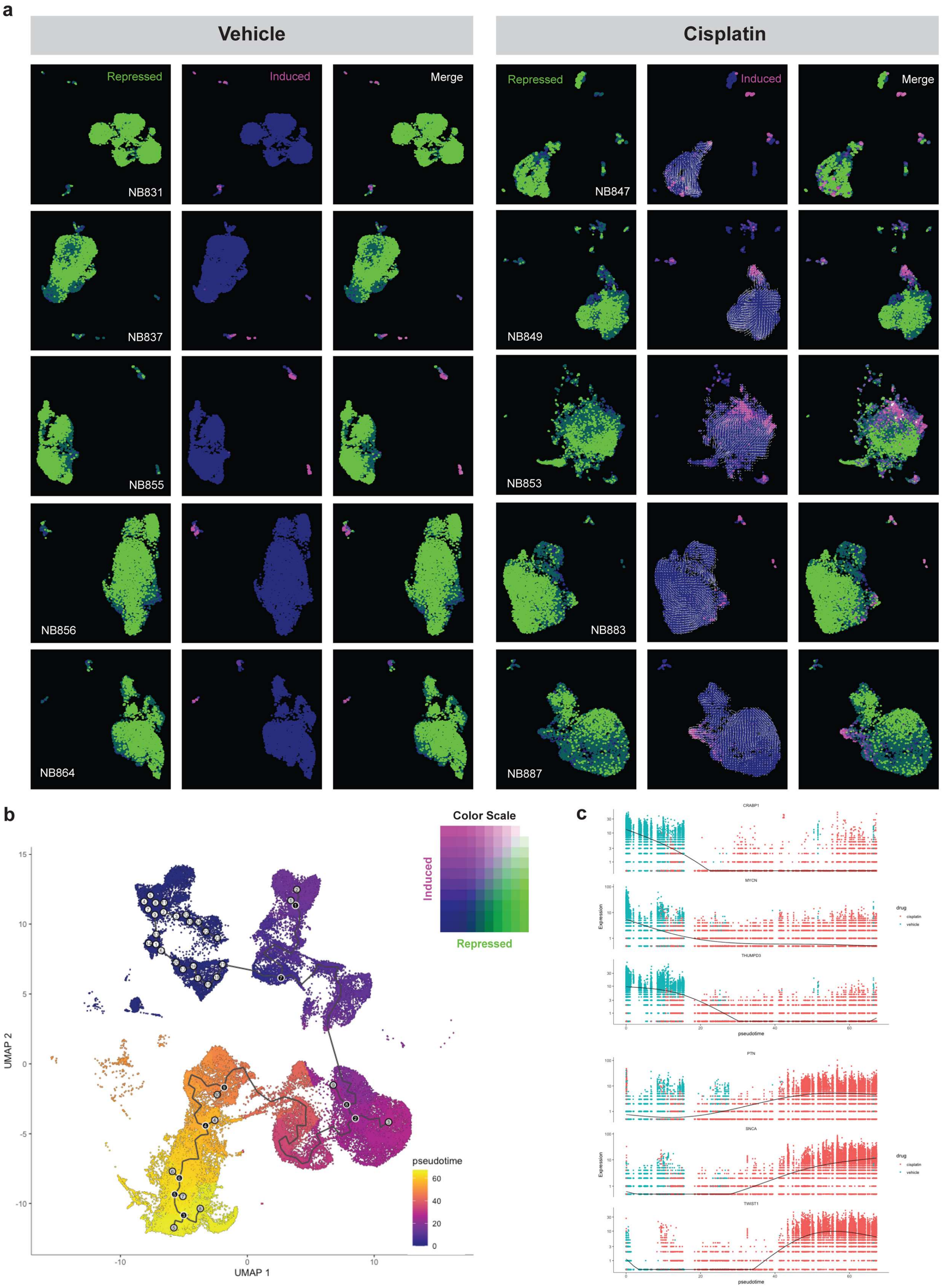
a
Vehicle
Cisplatin
Repressed
Induced
Merge
NB831
NB837
NB855
NB856
NB864
NB847
NB849
NB853
NB883
NB887
b
Color Scale
Induced
Repressed
UMAP 1
UMAP 2
pseudotime
c
CRABP1
MYCN
THUMPD3
PTN
SNCA
TWIST1
Expression
pseudotime
drug
cisplatin
vehicle

**Supplementary Figure 11. Examining transcriptional dynamics of drug response with RNA velocity and pseudotime.**

(a) RNA velocity was estimated in each mouse using scVelo. Cells are colored by one of two gene sets – those genes found to be significantly repressed after cisplatin treatment (green), and those genes induced by the chemotherapeutic (magenta). Whereas the velocities projected on the 2D embedding (UMAP) in the drug treated mice are all directed towards the cells most highly enriched for the induced gene set, the origin of these velocities arises from those cells most enriched in those genes that are repressed due to the treatment. Importantly, this was observed in every drug-treated mouse tumor. As the duration of this treatment was short (only 24 hours), the observation of these multiple cell states co-existing in these tumors likely represents an initiation of transcriptional changes with groups of cells existing in various stages of the response. Thus, the RNA velocity trajectory embeddings recover an accurate representation of the plausible cell state transitions.

(b) Monocle3 was used to derive a trajectory graph in single cell RNA-seq data from our GEMM. The root nodes were manually defined in untreated mouse samples (NB856 and NB864, specifically), and cells were plotted in pseudotime.

(c) Morans' I spatial correlation test was implemented on the principal graph of the trajectory inference in (b). The top three panels illustrate genes identified in our DESeq2 differential expression analysis that also co-varied in pseudotime (*Crabp1*, *MYCN*, and *Thumpd3*, respectively). The bottom panels show the reverse trend and are significantly increased in those cells at later pseudotime (*Ptn*, *Snca*, and *Twist1*, respectively). Each dot represents a cell from (b) and is colored by drug treatment.
